# Human skeletal muscle ageing atlas

**DOI:** 10.1101/2022.05.24.493094

**Authors:** Veronika R. Kedlian, Yaning Wang, Tianliang Liu, Xiaoping Chen, Liam Bolt, Zhuojian Shen, Eirini S. Fasouli, Elena Prigmore, Vitalii Kleshchevnikov, Tong Li, John E Lawrence, Ni Huang, Qin Guo, Lu Yang, Krzysztof Polański, Monika Dabrowska, Catherine Tudor, Xiaobo Li, Omer Bayraktar, Minal Patel, Kerstin B. Meyer, Natsuhiko Kumasaka, Krishnaa T. Mahbubani, Andy Peng Xiang, Kourosh Saeb-Parsy, Sarah A Teichmann, Hongbo Zhang

**Affiliations:** Wellcome Sanger Institute, Wellcome Genome Campus, Hinxton, Cambridge CB10 1SA, UK; Key Laboratory for Stem Cells and Tissue Engineering, Ministry of Education, Zhongshan School of Medicine, Sun Yat-sen University, Guangzhou, China; Advanced Medical Technology Center, The First Affiliated Hospital, Zhongshan School of Medicine, Sun Yat-sen University, Guangzhou, China; Department of Thoracic Surgery, Guangdong Provincial Key Laboratory of Malignant Tumor Epigenetics and Gene Regulation, Sun Yat-sen Memorial Hospital of Sun Yat-sen University, Guangzhou, China; Core Facilities for Medical Science, Sun Yat-sen University, Guangzhou, China; Department of Surgery, University of Cambridge, NIHR Cambridge Biomedical Research Centre, Cambridge Biorepository for Translational Medicine, Cambridge, UK; Cavendish Laboratory, University of Cambridge, JJ Thomson Ave, Cambridge CB3 0HE, UK

**Keywords:** Skeletal muscle, Ageing, MuSC, Myofiber, Single-cell RNA sequencing, Single-nucleus RNA sequencing

## Abstract

Skeletal muscle ageing increases the incidence of age-associated frailty and sarcopenia in the elderly worldwide, leading to increased morbidity and mortality. However, our understanding of the cellular and molecular mechanisms of muscle ageing is still far from complete. Here, we generate a single-cell and single-nucleus transcriptomic atlas of skeletal muscle ageing from 15 donors across the adult human lifespan, accompanied by myofiber typing using imaging. Our atlas reveals ageing mechanisms acting across different compartments of the muscle, including muscle stem cells (MuSCs), myofibers and the muscle microenvironment. Firstly, we uncover two mechanisms driving MuSC ageing, namely a decrease in ribosome biogenesis and an increase in inflammation. Secondly, we identify a set of nuclei populations explaining the preferential degeneration of the fast-twitch myofibers and suggest two mechanisms acting to compensate for their loss. Importantly, we identify a neuromuscular junction accessory population, which helps myofiber to compensate for aged-related denervation. Thirdly, we reveal multiple microenvironment cell types contributing to the inflammatory milieu of ageing muscle by producing cytokines and chemokines to attract immune cells. Finally, we provide a comparable mouse muscle ageing atlas and further investigate conserved and specific ageing hallmarks across species. In summary, we present a comprehensive human skeletal muscle ageing resource by combining different data modalities, which significantly expands our understanding of muscle biology and ageing.

## Introduction

Skeletal muscle makes up 40% of our body mass, and is not only an essential part of the locomotor system but also plays pivotal roles in metabolism and immune regulation^1–5^. The major components of skeletal muscle are multinucleated tubes called myofibers, which drive muscle contraction. Human myofibers are traditionally classified into “slow-twitch” (type I) and “fast-twitch” (type IIA and type IIX) according to their contraction speed, structural protein compositions and metabolic characteristics (more oxidative type I *vs.* more glycolytic type II). Myofibers are surrounded by mononuclear muscle stem cells (MuSCs) which can give rise to new myofibers and restore existing myofibers in the event of damage. In addition, there is a muscle microenvironment consisting of supporting fibroblasts, vasculature and immune cells, as well as Schwann cells and neuronal axons which transmit action potentials to the myofibers.

Skeletal muscle ageing is manifested by the loss of both muscle mass and strength, often leading to sarcopenia^6, 7^. This is a major contributory factor to falls and fractures in the elderly^8^, which globally represents the second leading cause of injury and deaths^9^. Myofiber ageing is associated with the selective decrease in both number and size of fast-twitch myofibers^10^. Furthermore, the number of MuSCs and their activation and proliferation in response to stimuli decrease with age, especially for MuSCs accompanying fast-twitch myofibers^10, 11^. However, it is not known whether this differential atrophy is due to myofiber-intrinsic changes in gene expression, the impact of the cellular microenvironment, or a combination of both. Several other putative muscle ageing factors, such as stem cell senescence, alterations in protein turnover, metabolic dysregulation, and chronic inflammation have been investigated in animal models^12–16^.

To date, most studies have focused on one particular mechanism or cell type in the muscle, and lacked a comprehensive approach to fully understand muscle ageing. To address this, recent mouse and human skeletal muscle studies pioneered the use of either single-cell (scRNA-seq)^17–21^ or single-nucleus RNA sequencing (snRNA-seq)^22–26^ to understand muscle cell type heterogeneity as well as their changes in ageing. Both sc- and snRNA-seq approaches have their own limitations when applied to muscle: droplet single-cell sequencing approaches are not well suited for myofiber profiling due to the multinuclearity and large size of fibres, while single-nucleus sequencing often lacks resolution for the less-abundant MuSCs and microenvironment cell types.

To obtain a systemic view of skeletal muscle ageing, we performed matched scRNA-seq and snRNA-seq of intercostal muscle across the adult human lifespan. This allowed us to investigate transcriptional changes of MuSCs, myofibers and microenvironment cells during ageing. We highlighted a set of interactions between these cells that could contribute to the ageing phenotype. We also performed myofiber typing of the intercostal muscle, providing a way to connect traditional observations about myofiber dynamics with changes occurring at the single-nucleus level. Finally, we generated age-matched single-cell and single-nucleus transcriptomes from mouse muscle, thus enabling the comparison of ageing across species.

## Results

### A single-cell and single-nucleus transcriptomic atlas of human skeletal muscle ageing

To gain a comprehensive view of human skeletal muscle ageing, we profiled the transcriptomes of 89,989 cells and 95,847 nuclei from intercostal muscle biopsies of 8 young (approx. 20-40 years old) and 7 aged donors (approx. 60-75 years old, Fig. 1a, b, Supplementary Table 1). Isolated cells and nuclei from each sample were subjected to droplet-based 3’ end sequencing (10x Genomics, Fig. 1a, Extended Data Fig. 1a-c).

**Fig. 1.**
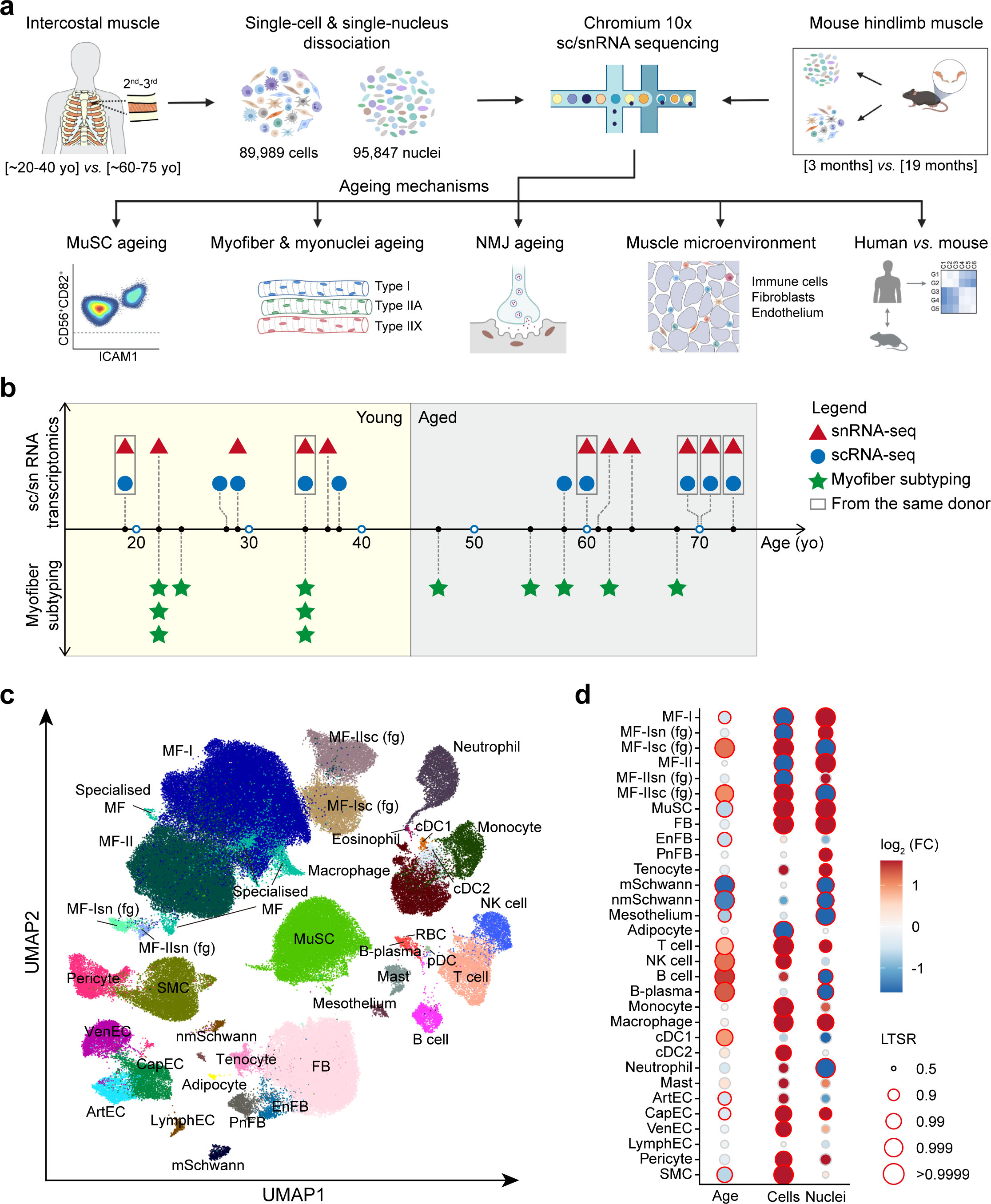
A single-cell and single-nucleus transcriptomic atlas of human skeletal muscle ageing. **a**, Visual overview of experimental design and main directions of investigations. Human intercostal muscles were collected between the 2^nd^ to the 3^rd^ rib, while mouse muscle samples were obtained from the whole hindlimbs. **b**, Timescale displaying human muscle sampling across ages for sc/snRNA-seq (n = 8 for young *vs.* n = 7 for aged) and for myofiber subtyping (n = 7 for young *vs.* n = 5 for aged). yo, years old. **c**, Uniform manifold approximation and projection (UMAP) visualisation of annotated cells in muscle ageing cell atlas. MF-I and MF-II, type I and type II myofiber; MF-Isn (fg) and MF-IIsn (fg), type I and type II myofiber fragment from snRNA-seq; MF-Isc (fg) and MF-IIsc (fg), type I and type II myofiber fragment from scRNA-seq; Specialised MF, specialised myonuclei (NMJ, NMJ accessory, MTJ) and myocyte (RASA4^+^, MYH8^+^) populations (to be described at Fig. 3); MuSC, muscle stem cell; FB, fibroblast; EnFB, endoneurial fibroblast; PnFB, perineurial fibroblast; mSchwann, myelinating Schwann cell; nmSchwann, non-myelinating Schwann cell; B-plasma, plasma cell; cDC1 and cDC2, conventional dendritic cell 1 and 2; ArtEC, arterial endothelial cell; VenEC, venous endothelial cell; CapEC, capillary endothelial cell; LymphEC, lymphatic endothelial cell; SMC, smooth muscle cell. **d**, The log_2_ (Fold change, FC) in the abundance of cell clusters across age (first column), and enrichment in cells compared to nuclei fraction (second and third columns), taking into account confounding factors (Methods). LTSR represents a significance measure indicating the local true sign rate. Significantly differentially abundant populations are highlighted with red edges (Methods).

After batch correction and integration of single-cell and single-nucleus data with the single-cell variational inference (scVI) autoencoder^27^, we annotated 39 major human cell populations, each displaying canonical marker genes (Fig. 1c, Extended Data Fig. 1a, Supplementary Table 1). We identified mononucleated MuSCs (*PAX7*^+^ and *DLK1*^+^), fibroblasts (FBs, *DCN*^+^), smooth muscle cells (SMCs, *ACTA2*^+^ and *MYH11*^+^), pericytes (*KCNJ8*^+^ and *RGS5*^+^), endothelial cells (ECs, *PECAM1*^+^), adipocytes (*ADIPOQ*^+^ and *PLIN1*^+^), myelinating (mSchwann, *MPZ*^+^ and *MBP*^+^) and non-myelinating (nmSchwann, *L1CAM1*^+^ and *CDH19*^+^) Schwann cells, immune cells (*PTPRC*^+^), and finally the multinucleated myofibers (MF, *TNNT1*^+^ and *TNNT3*^+^). All cell populations were identified across age groups (Extended Data Fig. 1b).

While for most of the cell types, data were generated by both cellular and nuclear RNA sequencing, we used a mixed-effect Poisson regression model to assess preferential enrichment of cell types by each technology (Fig. 1d, Extended Data Fig. 1c, see Methods). This analysis indicated that SMCs, arterial (ArtEC) and venous ECs (VenEC), Schwann and most of the immune cells were mainly captured from scRNA-seq. The two techniques captured comparable populations for MuSCs and fibroblasts. In contrast, myofiber and adipocytes were almost entirely captured from nuclei sequencing (Extended Data Fig. 1c). Together, this illustrates the advantages of combining both techniques.

Next, we examined changes in cell type composition with age using the same mixed effect Poisson regression model (see Methods). Muscle samples from aged donors were strongly enriched for immune cells, including B cells (B cell, B-plasma), natural killer cells (NK cell), T cells (T cell), dendritic cells (cDC1) as well as myofiber fragments (MF-Isc (fg) and MF-IIsc (fg)) detected in scRNA-seq (Fig. 1d). The increase in B and T cells is consistent with studies in the ageing mouse brain, liver and adipose tissue^28, 29^ and provides evidence of immune infiltration in ageing muscle. Conversely, MuSCs, Schwann cells, endoneurial fibroblasts (EnFB), SMCs as well as arterial and capillary ECs (CapEC) were strongly depleted in aged individuals (Fig. 1d), consistent with the dysfunction of physiological activities in skeletal muscle of aged individuals^1^.

To compare the process of muscle ageing between species, we generated a mouse muscle ageing atlas by sequencing 68,956 cells and 27,573 nuclei from hindlimb muscles of 5 young (3 months) and 3 aged (19 months) C57BL/6 mice (Extended Data Fig. 1d-g). This atlas yielded comparable major cell types to human skeletal muscle (Extended Data Fig. 1d-g, Supplementary Table 1), including MuSCs, SMCs, ECs, fibroblasts, adipocytes, Schwann cells, immune cells and different states of myonuclei, thus providing a data resource for capturing cross-species muscle ageing hallmarks.

Overall, our scRNA-seq and snRNA-seq datasets identified major cell types residing in skeletal muscle, generating a comprehensive atlas, which we provide as an online resource for easy browsing and data download https://www.muscleageingcellatlas.org.

### Mechanistic insights into human MuSC ageing

In several animal models, the regenerative capacity of skeletal muscle declines with age, largely due to the senescence of MuSCs^30–33^. However, the extent of MuSC functional decline in human ageing remains a controversial topic^34, 35^. To better understand their age-related transcriptional changes, we performed unsupervised clustering of the 15,606 high-quality MuSCs across ages from the scRNA-seq data subset and identified 4 subpopulations (Fig. 2a, Supplementary Table 2). As expected, one of these populations represented the main type of MuSCs which expressed pan-MuSC lineage markers *PAX7* and *SPRY1*, demonstrating a quiescent state. The second identified state is the MYOG^+^ (MyG^+^) MuSC, that was highly enriched for both myogenic differentiation regulators *MYOD1* and *MYOG* as well as myofiber structural proteins such as *TNNT1* and *TNNT2*, representing a transient differentiating state (Fig. 2a, right panel). Intriguingly, we identified two additional subpopulations which we refer to as TNFRSF12A^+^ (TNF^+^) and ICAM1^+^ (ICA^+^) MuSC, respectively. Gene Ontology (GO) enrichment analysis of their differentially expressed genes (DEGs, Supplementary Table 2) indicated that several pathways enriched in the TNF^+^ population were related to development (Fig. 2b). Notably, ribosome biogenesis, which is required for MuSC activation and proliferation^36^, was the top enriched pathway in the TNF^+^ subpopulation (Fig. 2b, Supplementary Table 2). This points to TNF^+^ subpopulation representing activated MuSC, which plays a role in muscle regeneration. Indeed, the human TNF^+^ population shared numerous DEGs with murine activated MuSC (Extended Data Fig. 2a)^19,37–40^.

**Fig. 2.**
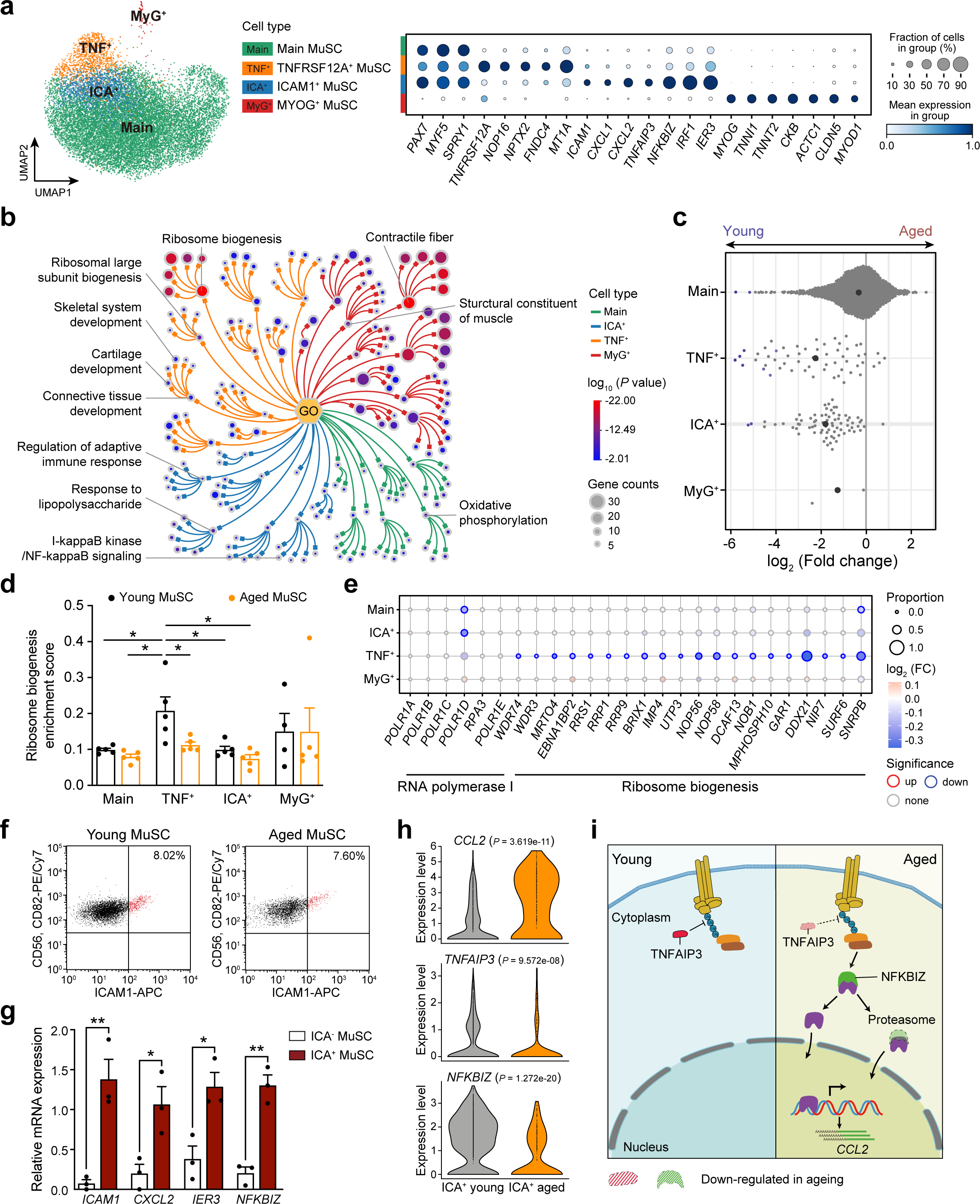
Mechanistic insights into human MuSC ageing. **a**, UMAP visualisation of 15,606 MuSCs, displaying main subpopulations identified from scRNA-seq (left) using marker genes from the dot plot (right). Size of the dot represents the proportion of cells expressing each gene, colour denotes the mean scaled expression level in each cluster. **b**, Tree visualisation of the clusters of Gene Ontology (GO) terms enriched among marker genes for every MuSC subpopulation (denoted with the colour of the branches) using Metascape (see Methods). Nodes represent GO terms, which are grouped into clusters based on semantic similarity (only top 10 clusters per each MuSC population and top 5 enriched terms within each cluster are shown). The most enriched GO term within a cluster is also chosen as a parent node, all parent nodes are connected to one arbitrary root. Size of the node denotes GO term size. Colour denotes log10 of P value. c, Beeswarm Milo plot (see Methods) showing the distribution of log_2_-fold change (x axis) in cell abundance with age across neighbourhoods of MuSC subtypes (y axis). Neighbourhoods displaying significant differential abundance at SpatialFDR 10% are coloured. **d**, Bar plot visualisation of ribosome biogenesis enrichment score among four MuSC subpopulations in young *vs.* aged human individuals. *P* values were obtained using a one-way ANOVA test. *, *P* < 0.05; **, *P* < 0.01. **e**, Dot plot illustrating ageing changes in the ribosome biogenesis genes and genes comprising RNA polymerase I complexes in each of the four MuSC subpopulations. Size of the dot represents the proportion of aged cells expressing the gene, colour denotes log_2_ (FC) in aged *vs*. young gene expression. Significantly up-regulated genes are highlighted with red and down-regulated with blue edges. **f**, **g**, FACS-based scatter plots (n = 4 for young *vs.* n = 4 for aged) of ICA^+^ MuSC (**f**) and quantitative real-time PCR (n = 3) validations based on its marker genes including *ICAM1*, *CXCL2*, *IER3* and *NFKBIZ*. (**g**). *P* values were obtained using unpaired two-tailed *t*-test. *, *P* < 0.05; **, *P* < 0.01. **h**, Violin plots display distribution of gene expression levels and their ageing changes for *CCL2*, *TNFAIP3* and *NFKBIZ* in ICA^+^ MuSC population from scRNA-seq data. **i**, Schematic diagram explaining the pro-inflammatory mechanism mediated by elevated NF-kB-CCL2 signalling pathway in ICA^+^ MuSC ageing. All data presented in (**d**, **g**) are expressed as mean ± SE with individual data points shown.

We then examined changes in cell abundance across conditions using a framework for differential abundance testing (Milo)^41^. Upon ageing, all four MuSC subpopulations decreased in number (Fig. 2c), which is consistent with most of the reports suggesting that MuSC function declines with age. The general decline in cell abundance was further experimentally validated using fluorescence-activated cell sorting (FACS) enriched CD31^-^CD82^+^CD56^+^ MuSCs^42, 43^ (Extended Data Fig. 2b, c). Interestingly, TNF^+^ population showed the greatest decrease in abundance among all four MuSC groups (Fig. 2c), suggesting a decrease in MuSC activation in ageing. The decline in the activation is one of the hallmarks of MuSC senescence in rodent models^30^. Our data, therefore, indicate a similar process occurs in humans during ageing. For instance, we found that the ribosome biogenesis genes and *POLR1D*, encoding a key subunit involved in the formation of the assembly platform for RNA polymerase I, both showed a tendency to decrease in most of the aged MuSC populations (Fig. 2d, e, Supplementary Table 3). Decrease of RNA polymerase I and ribosome biogenesis has been shown in several models to be a leading cause of stem cell senescence^36, 44, 45^. Importantly, the TNF^+^ population exhibited the greatest decrease in ribosome biogenesis enrichment score among all groups (Fig. 2d, Supplementary Table 2). In summary, our data demonstrate senescence of human MuSCs in ageing as a result of perturbation of ribosome biogenesis.

Next, we found that another subtype of MuSCs, namely ICA+ MuSC, showed enrichment in cytokine genes including *CXCL1*, *CXCL2*, *IRF1*, *IER3* and relevant NF-kB regulators *TNFAIP3* and *NFKBIZ* (Fig. 2a, Extended Data Fig. 2d), indicating strong immune-responsive properties. Using ICAM1 as a cell sorting marker (Fig. 2f), we confirmed the specific expression of *CXCL2*, *IER3* and *NFKBIZ* in ICA^+^ MuSCs (Fig. 2g). Immunoregulation is indispensable for successful MuSC regeneration^46^, whereas ageing is usually accompanied by chronic inflammation in the MuSC niche, reflecting dysregulated immune homeostasis upon MuSC senescence^1, 47^. Cell sorting also confirmed the tendency of ICA^+^ population to decline in ageing (on average 8.7% *vs.* 6.4% of total MuSC in young and aged individuals, respectively) (Extended Data Fig. 2e). More importantly, we found that upon ageing, the proinflammatory cytokine gene *CCL2* was dramatically up-regulated in ICA^+^ MuSCs (Fig. 2h, Supplementary Table 3), which was also confirmed by quantitative real-time PCR (Extended Data Fig. 2f). CCL2 can be tightly regulated by *NFKB1* transcription factor^48^. Indeed, two classical *NFKB1* inhibitor genes, *TNFAIP3* and *NFKBIZ*, were markedly reduced in ageing (Fig. 2h, Supplementary Table 3), indicating increased activity of NF-κB signalling. Thus, elevated activation of NF-κB signalling in ageing ICA^+^ MuSCs could lead to increased *CCL2* transcription (Fig. 2i), which contributes to impaired immune homeostasis and chronic inflammation in the muscle stem cell niche.

### Myofiber type-specific ageing mechanisms

Aiming to profile the human myofiber and the heterogeneity of its nuclei in young and aged skeletal muscle, we isolated myonuclei data from both sc- and snRNA-seq and complemented this data with immunofluorescence-based myofiber typing performed on the tissue sections (Fig. 3a).

**Fig. 3.**
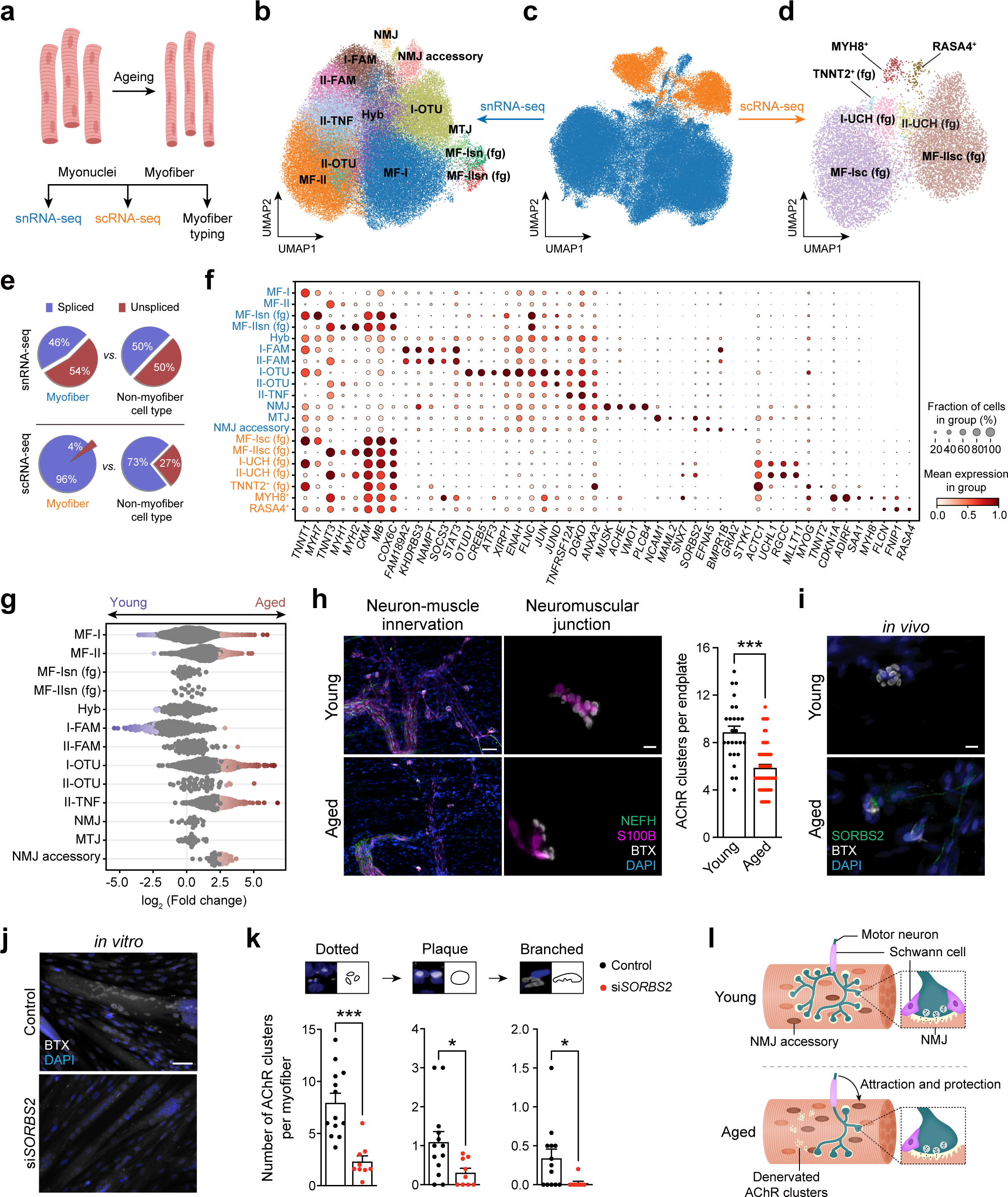
Myofiber type-specific ageing mechanisms. **a**, A schematic diagram illustrating myofiber ageing and main approaches used for its investigations. **b**-**d**, UMAP visualisation of myofiber populations obtained from integrated sn- and scRNA-seq (**c**) as well as separate visualisations of populations recovered from sn- (**b**) and scRNA-seq (**d**). **e**, Pie charts illustrating the difference in the average ratio of spliced and unspliced transcripts in myofiber nuclei (from **b**) and cell populations (from **d**) as opposed to nuclei and cells comprising other cell types (immune, stromal cells and others). **f**, Dot plot showing marker genes for myonuclei and cell populations derived from sn- (blue) and scRNA-seq (orange) data. Size of the dot represents the proportion of cells expressing a gene. Colour denotes scaled expression level. **g**, Beeswarm Milo plot (see Methods) showing the distribution of log_2_-fold change (x axis) in cell abundance with age across neighbourhoods of myonuclei populations (y axis). Neighbourhoods displaying significant differential abundance at SpatialFDR 10% are coloured. **h**, Immunofluorescence (IF, left panels) on teased human intercostal muscles shows neuromuscular junction architecture and its morphological changes in ageing. Acetylcholine receptors (AChRs) were stained with BTX (white), nuclei with DAPI (blue), axons with anti-NEFH antibodies (green) and Schwann cells with anti-S100B antibodies (violet). Changes in AChR clusters upon ageing were also statistically analysed (right panel, n = 1 for young *vs.* n = 1 for aged). *P* value was obtained using unpaired two-tailed *t*-test. Scale bar (left): 50 µm; Scale bar (right): 10 µm. ***, *P* < 0.001. **i**, IF co-staining of AChRs with BTX (white) and NMJ accessory nuclei with anti-SORBS2 antibodies (green) in young and aged human intercostal muscles. Scale bar: 10 µm. **j**, **k**, IF staining of AChRs with BTX (white) in cultured human myotubes before and after siRNA-mediated knockdown of *SORBS2* expression (**j**). Bar plots showing the number of AChR clusters per myotube for three different stages of cluster formation (Dotted to Plaque to Branched) before and after *SORBS2* knockdown. Images were analysed by Fiji (**k**, n = 15 fields for control *vs.* n = 9 fields for si*SORBS2* group). *P* values were obtained using unpaired two-tailed *t*-test with (Plaque and Branched) or without (Dotted) Welch’s correction. Scale bar: 50 µm. *, *P* < 0.05; ***, *P* < 0.001. **l**, Schematic diagram showing NMJ accessory nuclei-mediated prosurvival mechanism against NMJ ageing. All data presented in bar plots (**h**, **k**) are expressed as mean ± SE with individual data points shown.

By integrating scRNA-seq and snRNA-seq myofiber data using the scVI model, we obtained 90,106 myofiber cells/nuclei (Fig. 3c, Extended Data Fig. 3a), clustering into 6 major populations and a few specialised populations. Myofiber types were then annotated according to conventional markers: *MYH7, TNNT1, TNNC1* for slow-twitch myofiber; and *MYH1, MYH2, TNNT3, TNNC2* for fast-twitch myofiber. Among the 6 main populations, two pairs of slow-twitch (myofiber type I, MF-I and MF-Isn (fg)) and fast-twitch myofibers (MF-II and MF-IIsn (fg)) came from single-nucleus data, and one pair of MF-Isc (fg) and MF-IIsc (fg) originated from single-cell data (fg denotes myofiber fragment). Interestingly, scRNA-seq-derived and snRNA-seq-derived fragment populations (MF-Isc (fg), MF-IIsc (fg), MF-Isn (fg), MF-IIsn (fg)) contained a much larger percentage of spliced transcripts (>= 97% and 92%, respectively, Extended Data Fig. 3b) in comparison with cells and nuclei derived from non-myofiber cell types (73% and 50%, respectively) (Fig. 3e). We hypothesised that the increased spliced/unspliced ratio in these four states was a result of myofiber partitioning or debris contamination, often occurring during the isolation process. This was also supported by the high expression of abundant sarcoplasmic and mitochondrial transcripts such as *CKM*, *MB*, *COX6C*, *ATP5H* in the fragment populations (Fig. 3f). We, therefore, focused on the populations with lower percentage of spliced transcripts, which likely represented pure myofiber cells or nuclei.

Further subclustering of myofiber single-cell and single-nucleus data revealed 9 additional nuclei and 5 cell populations (Fig. 3b, d, f, Supplementary Table 4). From snRNA-seq, we identified paired FAM189A2^+^ nuclei (I-FAM, II-FAM) and OTUD1^+^ nuclei (I-OTU, II-OTU) representing convergent states present in both slow-twitch and fast-twitch myofibers. We also observed a novel fast-twitch myofiber specific TNFRSF12A^+^ (II-TNF) and three specialised myonuclei states, which included previously described neuromuscular junction (NMJ) nuclei, myotendinous junction (MTJ) nuclei and a novel neuromuscular junction accessory state (NMJ accessory). In the single-cell dataset, we observed two putative myocyte populations (as inferred by the ratio of spliced transcripts, Extended Data Fig. 3c) that were marked by *MYH8* and *RASA4* and three myofiber fragment populations. Intriguingly, MYH8^+^ myocytes specifically expressed fetal myosin heavy chain *MYH8* in addition to *CDKN1A* and *MYOG*, indicating active myogenesis. In turn, RASA4^+^ myocytes can be involved in muscle growth given that they expressed FLCN/FNIP1 complex and the GTPase activating protein RASA4, both of which are activators of growth sensing regulator mTORC1^49–51^.

By investigating age-associated cell composition changes using the Milo tool, we uncovered that the dynamics of myonuclei states with age differed between myofiber types (Fig. 3g). First, FAM189A2^+^ myonuclei substantially decreased while OTUD1^+^ nuclei increased in slow-twitch myofibers with age. In contrast, these two populations did not change in fast-twitch myofibers, but instead an increase in the TNFRSF12A^+^ population was observed.

FAM189A2^+^ nuclei were characterised by the expression of *NAMPT*^52, 53^, an enzyme which promotes NAD^+^ replenishment, as well as *STAT3*, a transcription factor which is activated by Janus kinase-mediated phosphorylation in response to cytokines and growth factors^54^, and its negative regulator *SOCS3*^55^ (Fig. 3f). Activation of STAT3 signalling was reported to be required for muscle repair^56, 57^. However, high and sustained activation of the JAK-STAT pathway induces inflammation and muscle dystrophy, especially in aged rodents^58–60^. Additionally, FAM189A2^+^ nuclei showed enrichment in the stress response (Extended Data Fig. 3d, Supplementary Table 4). Taken together, we hypothesise that decreased FAM189A2^+^ nuclei population in slow-twitch myofibers is a result of a protective mechanism which reduces negative consequences of the stress response, at the cost of reducing the capacity for muscle repair with age.

OTUD1^+^ nuclei were characterised by expression of the deubiquitinating enzyme gene *OTUD1*, the injury activated transcription factor *CREB5* and its target *ATF3*, the three actin-cytoskeleton related genes *XIRP1*, *ENAH*, *FLNC,* and *DNAJA4*, a chaperone involved in the unfolded protein response (Fig. 3f). Indeed, OTUD1^+^ nuclei were enriched in active growth and repair supporting activities (Extended Data Fig. 3d, Supplementary Table 4). This agrees well with the fact that a population with similar marker genes was previously reported in mouse and implicated as “repair” cluster^22–24^. Therefore, increased OTUD1^+^ nuclei point to the activation of repair mechanisms preventing degeneration of slow-twitch myofibers.

TNFRSF12A^+^ nuclei had an expression profile similar to OTUD1^+^ state and showed specific enrichment in “wound healing” and “blood coagulation” gene sets, also indicating its muscle repair activity (Extended Data Fig. 3d, Supplementary Table 4). Besides, it also expressed higher levels of *TNFRSF12A* (Fig. 3f), which encodes FN14, a receptor for the TWEAK ligand that is well-known to promote muscle atrophy^61^. Hence, an increase in TNFRSF12A^+^ nuclei in fast-twitch myofibers can be a result of disproportional damage repair response which leads to the preferential atrophy of these myofibers^62, 63^. Taken together, FAM189A2^+^, OTUD1^+^ and TNFRSF12A^+^ nuclei may be responsible for differential susceptibility of slow-twitch and fast-twitch myofibers to age-related degeneration and atrophy.

Surprisingly, we also identified a novel nuclei population (named “NMJ accessory”), which showed enrichment in various neuronal GO terms such as “synapse organisation” and “axon development” as well as expressed marker genes related to synapse formation distinct from classical NMJ nuclei markers (Fig. 3f, Extended Data Fig. 3d, e, Supplementary Table 4). Among these markers were *GRIA2*, encoding the key subunit of ionotropic glutamate receptor AMPAR that mediates synaptic plasticity and promotes cholinergic transmission at NMJ^64, 65^, *EFNA5*, encoding an essential ligand involved in axon guidance to the myotube during limb development^66^, and *SORBS2*, encoding an adapter protein involved in acetylcholine receptor (AChR) cluster formation in mouse^67^. The “NMJ accessory” subpopulation also dramatically increased with age (Fig. 3g), suggesting its abundance may be responding to age-related changes in muscle innervation. Indeed, we found that the NMJ suffered severe degradation with age, displaying decreased AChR clusters and protective terminal Schwann cells (Fig. 3h). Using SORBS2 as a marker, we identified some NMJ accessory nuclei directly beneath the postsynaptic endplates at the NMJ of aged individual, but not the young individual (Fig. 3i). In further support of their functional importance, *SORBS2* knockdown in cultured human myotubes led to a dramatic decrease of AChR clusters on all stages of aggregate assembly (Fig. 3j, k). In summary, we report a novel population of nuclei which may compensate for degeneration and denervation of NMJ (Fig. 3l).

### Compensatory mechanisms against human fast-twitch myofiber degeneration in ageing

Myofibers have differential susceptibility to ageing depending on their type: fast-twitch myofibers are more vulnerable than slow-twitch ones^68^. Here we combined information about myofiber and nuclei dynamics to investigate this phenomenon. At the myofiber level, we used immunofluorescence staining of traditional myosin heavy chain proteins followed by image analysis (see Methods, Fig. 4a) to distinguish slow-twitch (type I, MYH7^+^), fast-twitch (type IIA, MYH2^+^ and type IIX, MYH1^+^) and hybrid (type IIA-IIX, MYH2^+^MYH1^+^) myofibers. We then scored the expression of the same genes in the nuclei, and consequently were able to separate three pure nuclei types, *MYH7*^+^, *MYH2*^+^, and *MYH1*^+^, as well as four hybrid nuclei types, *MYH2*^+^*MYH1*^+^, *MYH7*^+^*MYH1*^+^, *MYH7*^+^*MYH2*^+^ and *MYH7*^+^*MYH2*^+^*MYH1*^+.^ (Fig. 4a, Supplementary Table 5).

**Fig. 4.**
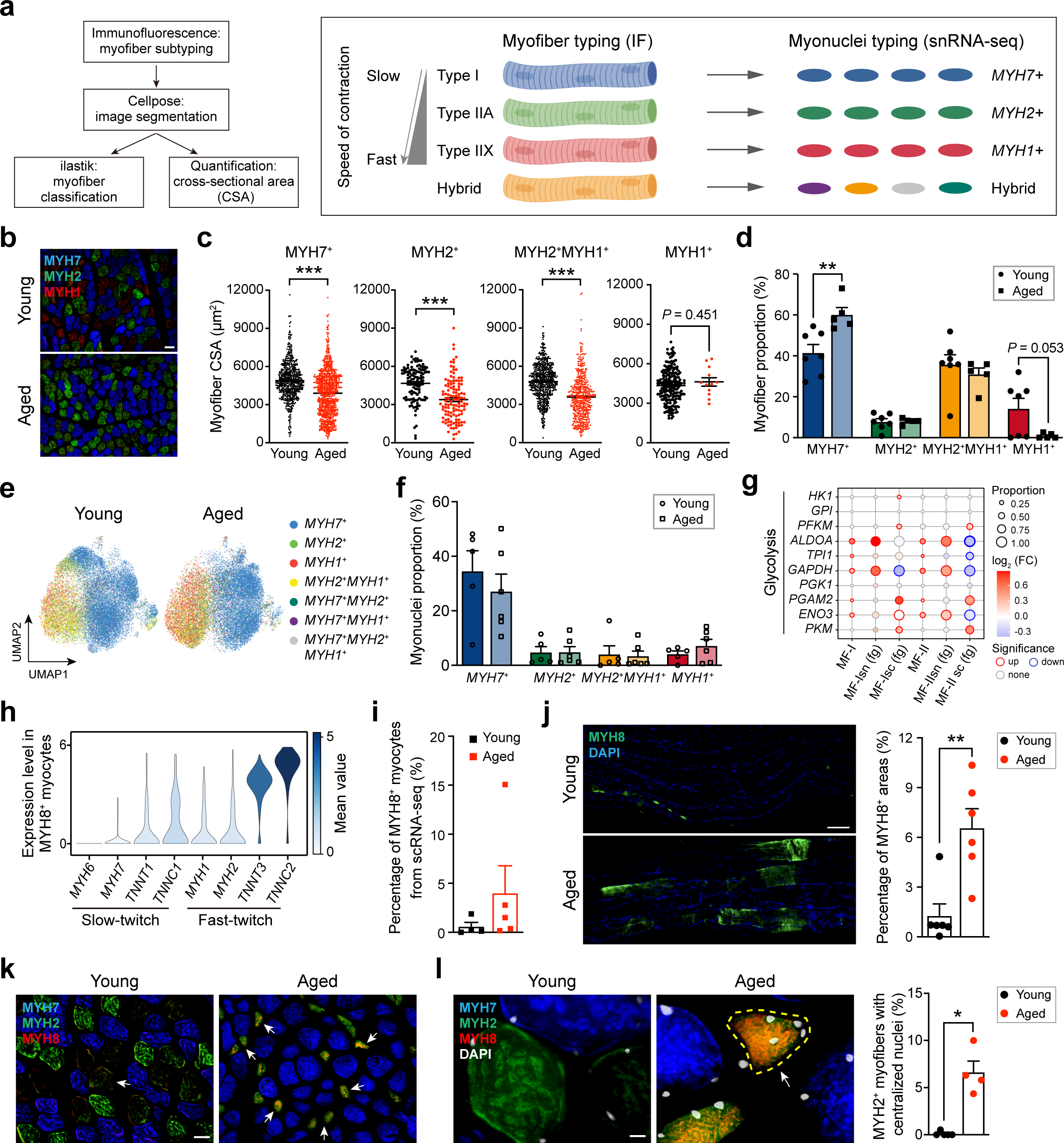
Compensatory mechanisms against human fast-twitch myofiber degeneration in ageing. **a**, Myofiber subtyping workflow and correspondence diagram between myofiber (typing done by immunohistochemistry (IHC)) and myonuclei types (typing done by snRNA-seq). **b**, IHC staining of different myofiber types in young and aged human individuals (n = 7 for young *vs.* n = 5 for aged). Anti-MYH7 antibody was used to stain type I (blue), anti-MYH2 to stain type IIA (green) and anti-MYH1 to stain type IIX (red) myofiber. Scale bar: 100 µm. **c**, Distribution of myofiber cross-sectional area (CSA, y axis) in young *vs.* aged (x axis) across different myofiber types. **d,** Paired bar plots showing the proportion of each myofiber type in young *vs.* aged individuals. Data were collected using automatic IHC image analysis (**c, d**). *P* values were obtained using unpaired two-tailed *t*-test with (**c**) or without (**d**) Welch’s correction. **, *P* < 0.01; ***, *P* < 0.001. **e**, UMAP visualisations of myofiber nuclei in young and aged individuals which were coloured according to myonuclei type. **f**, Paired bar plots showing the proportion of each myonuclei type in young *vs.* aged which were calculated based on data in (**e**). **g**, Dot plot illustrating age-related differential expression of glycolysis genes across six main myofiber populations. The size of the dot represents the proportion of aged cells expressing the gene. Colour denotes log_2_ (FC) in aged vs. young upon gene expression. Significant differentially expressed genes (DEGs) are highlighted with coloured edges. **h**, Violin plots generated from scRNA-seq data show the distribution of expression of slow-twitch and fast-twitch myofiber markers in MYH8^+^ myocytes. **i**, Bar plot showing the change in the proportion of MYH8^+^ myocytes, relative to the total myofiber cells obtained in scRNAseq, with age. **j**, IF on teased human intercostal muscles (left) showing expression of MYH8 (green) in young *vs.* aged myofibers. Bar plot illustrating corresponding percentage of MYH8^+^ areas (right) in young (n = 6) *vs.* aged (n = 6). *P* value was obtained using the unpaired two-tailed *t*-test. **, *P* < 0.01. Scale bar: 100 µm. **k**, **i**, IHC staining shows MYH8 expression (red) on skeletal muscle cross-sections in young vs aged in low (**k,** scale bar: 50 µm) and high-resolution (**i**, left panel, scale bar: 10 µm). Type I, type IIA and nuclei are stained with anti-MYH7 (blue), anti-MYH2 (green) antibodies and DAPI (white). Arrows point to MYH8^+^ myofibers. **l**, Bar plot (right panel) showing the corresponding proportion of MYH2^+^ myofibers with centralised nuclei in young (n = 5) *vs.* aged (n = 4) individuals. *P* value was obtained using unpaired two-tailed *t*-test with Welch’s correction. *, *P* < 0.05. All data in (**c**, **d**, **f**, **i**, **j**, **l**) are expressed as mean ± SE with individual data points shown.

At the myofiber level, we confirmed reduced heterogeneity of fast-twitch myofibers in aged compared to young intercostal muscle (Fig. 4b, Extended Data Fig. 4a). We also showed that both slow-twitch and fast-twitch myofibers decreased in cross-sectional area (CSA), with type IIA displaying the greatest reduction in size (Fig. 4c, Extended Data Fig. 4b-d). Detailed myofiber typing revealed that type IIX myofibers almost completely disappeared in aged individuals, indicating the strongest degeneration. Type IIA and hybrid IIA-IIX did not significantly change, while type I had increased in proportion (Fig. 4d, Extended Data Fig. 4e, Supplementary Table 5).

At the nuclei level, there were no significant changes between young and aged muscle (Fig. 4e, f, Extended Data Fig. 4f). However, *MYH1*^+^ nuclei (type IIX) had a tendency to increase, even though type IIX myofibers disappeared entirely with age (Fig. 4f). This may point to initiation of MYH1 expression in other myonuclear types (*MYH7*^+^, *MYH2*^+^) and acquisition of a hybrid nuclear phenotype. In agreement with this, expressions of typical glycolytic genes were dramatically increased in slow-twitch myonuclei (Fig. 4g), in keeping with a previous report^68^, while expressions of main mitochondrial biogenesis genes *PPARGC1A* and *NRF1* were decreased (Extended Data Fig. 4g). Such hybrid states may work to compensate for fast-twitch myofiber loss in ageing (Supplementary Table 5).

Meanwhile, MYH8^+^ myocytes (previously identified from scRNA-seq) may present another compensatory mechanism for fast-twitch myofiber loss. In particular, we found MYH8^+^ myocytes to be an intermediate state in the trajectory from MuSC to myofiber predominantly linking MuSC to fast-twitch myofiber (Extended Data Fig. 4h, i). This is consistent with neonatal MYH8^+^ being previously described as a marker of muscle regeneration^69^. MYH8^+^ myocytes also specifically expressed high levels of fast-twitch myofiber markers and increased in proportion with age (Fig. 4h, i, Supplementary Table 5). We validated the increase in MYH8 expression with age and its co-expression with fast-twitch myofiber markers on the tissue sections *in vivo* (Fig. 4j, k). Moreover, nearly 10% of MYH8^+^ fast-twitch myofibers had centralised nuclei (Fig. 4l), suggesting they had recently initiated regeneration.

In summary, taking advantage of immunofluorescence myofiber typing together with sn/scRNA-seq, our data reveal two novel compensatory mechanisms for fast-twitch myofiber loss: oxidative-to-glycolytic shift in nuclei states and an increase in fast-twitch myofiber regeneration via MYH8^+^ myocytes.

### Cell type composition of human skeletal muscle microenvironment and ageing regulation

The muscle microenvironment has been proposed to play an indispensable role in muscle ageing^70^. To address this claim, we separated and finely annotated major populations comprising both myofiber and MuSC microenvironments including immune cells, fibroblasts, Schwann cells, ECs and SMCs (Fig. 5a-c, Extended Data Fig. 5a-c, Supplementary Table 6).

**Fig. 5.**
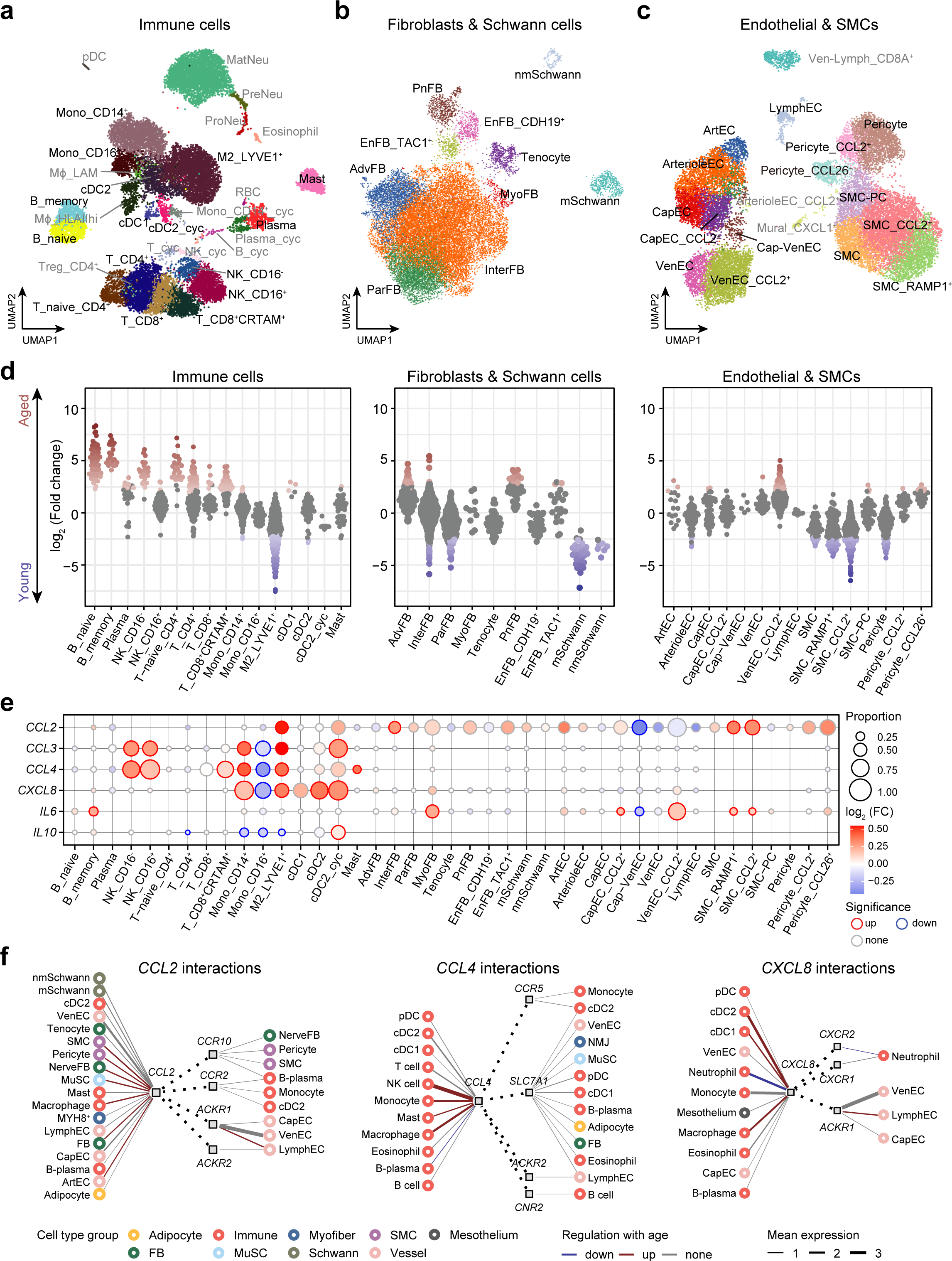
Cell type composition of human skeletal muscle microenvironment and ageing regulation. **a**-**c**, UMAP plots showing annotated subpopulations of immune cells (**a**), fibroblasts and Schwann cells (**b**), as well as endothelial and smooth muscle cells (**c**). Cycling cells: T_cyc, NK_cyc, B_cyc, Plasma_cyc, Mono_CD14^+^_cyc, cDC2_cyc; Mono_CD14^+^, CD14^+^ monocytes; Mono_CD16^+^, CD16^+^ monocytes; M2_LYVE1^+^, LYVE1 M2 macrophages; Mϕ_HLAIIhi, MHCII high macrophages; Mϕ_LAM, lipid-associated macrophages; cDC1 and cDC2, conventional dendritic cell 1 and 2; Mast, mast cell; pDC, plasmacytoid dendritic cell; ProNeu, pro-neutrophil; PreNeu, pre-neutrophil; MatNeu, mature neutrophil; AdvFB, adventitial fibroblasts; InterFB, intermediate fibroblasts; ParFB, parenchymal fibroblasts; MyoFB, myofibroblasts; EnFB_TAC1^+^ and EnFB_CDH19^+^, endoneurial fibroblast TAC1^+^ and CDH19^+^; PnFB, perineurial fibroblast; mSchwann, myelinating Schwann cell; nmSchwann, non-myelinating Schwann cell; ArtEC, arterial endothelial cell; ArterioleEC, arteriole endothelial cell; VenEC, venous endothelial cell; CapEC, capillary endothelial cell; Cap-VenEC, capillary venous endothelial cell; LymphEC, lymphatic endothelial cell; SMC, smooth muscle cell; SMC-PC, smooth muscle cell/pericyte cell. Cell populations marked in grey contained very few cells or (and) were represented in 1-2 individuals, thus were excluded from further analyses. **d**, Beeswarm Milo plot (see Methods) showing the distribution of log_2_-fold change (y axis) in cell abundance with age across neighbourhoods of immune cells (**a**), fibroblasts and Schwann cells (**b**), endothelial and smooth muscle cell subtypes (**c**) (y axis). Neighbourhoods displaying significant differential abundance at SpatialFDR 10% are coloured. **e**, Dot plot illustrating ageing changes in the set of chemokines and interleukins among muscle environment subpopulations. Size of the dot represents the proportion of aged cells expressing the gene. Colour denotes log_2_ (FC) in aged *vs.* young upon gene expression. Significantly up-regulated genes are highlighted with red and down-regulated with blue edges. **f**, Putative cell-cell interactions in the aged skeletal muscle mediated via *CCL2*, *CCL4* and *CXCL8* chemokines, produced by microenvironment cells. Emitter (leftmost) and receiver (rightmost) cell types are marked with circles, which are colored according to a broad cell type group; ligands and receptors are marked with square nodes. Solid edges connect cell types and ligands, or receptors, which they express; thickness of the line is proportional to the mean expression level of the gene in cell type. Significantly differentially expressed genes (Supplementary Table 3) are coloured. Dotted edges connect putative receptors and their ligands.

In the immune cell compartment (Fig. 5a, Extended Data Fig. 5a, Supplementary Table 6), we identified various subtypes of lymphoid and myeloid cells including T cells (CD4^+^, CD8^+^, CD8^+^CRTAM^+^^71^), B cells (naive and memory), plasma cells, NK cells (CD16^+^, CD16^-^), monocytes (Mono, CD14^+^, CD16^+^), dendritic cells (cDC1, cDC2 and pDC) and macrophages (M2_LYVE1^+^^72–74^, Mϕ_HLAIIhi^75^ and Mϕ_LAM^74, 76^) which have been reported by recent studies across a range of tissues. In two donors we also detected granulocytes, specifically neutrophils at various stages of development, and eosinophils, which play a role in muscle regeneration^77^.

In the stroma-neural compartment (Fig. 5b, Extended Data Fig. 5b, Supplementary Table 6), we observed tenocytes (*TNMD*^+^ and *SCX*^+^), a large population of adventitial fibroblasts (AdvFB, *PI16*^+^), a population of parenchymal fibroblasts (ParFB, *COL4A1*^+^ and *COl15A1*^+^) and a population of intermediate fibroblasts (InterFB)^78^, a transitional state between AdvFB and ParFB. We also identified mSchwann- and nmSchwann cells, some of which displayed markers of terminal Schwann cells^79^ (Extended Data Fig. 5d), known to specifically localise at and protect the NMJ. In addition, we noted a number of nerve-associated fibroblasts (NerveFB)^80, 81^, including perineurial (PnFB) and endoneurial (EnFB) subtypes. We described a novel subtype of endoneurial fibroblasts expressing neuropeptide Tachykinin Precursor 1 (*TAC1*), known to exert a wide range of effects on nerve and smooth muscle cells.

In the vascular compartment (Fig. 5c, Extended Data Fig. 5c, Supplementary Table 6), we identified main endothelial cell types such as arteria (ArtEC), vein (VenEC), capillaries (CapEC), lymphatics (LymphEC) together with the pericytes, SMCs and mural cells that form blood vessel walls. Across the blood vessel cell types, we observed convergent inflammatory response states, in both young and aged, characterised by expression of interferon regulatory factor, *IRF1, CCL2* cytokine as well as DNA damage response genes *GADD45B, CDKN1A*.

By investigating age-associated dynamics of microenvironment populations using Milo (Fig. 5d), we discovered that the largest increase in abundance with age occurred among the lymphoid immune cells including B cells, T cells, NK_CD16^-^ cells, as well as with certain fibroblast subtypes (AdvFB and PnFB). Within the blood vessel compartment, CCL2^+^ veins (VenEC_CCL2^+^) displayed the strongest enrichment in elderly donors and were followed by CCL2^+^ (Pericyte_CCL2^+^) and CCL26^+^ pericytes (Pericyte_CCL26^+^). These findings add insight to previous observations that the incidence of diseases involving muscle inflammation and fibrosis increases with age^82, 83^. In contrast, both mSchwann- and nmSchwann cells dramatically decreased with age, in keeping with the accompanying axonal deterioration. Interestingly, anti-inflammatory M2-like LYVE1^+^ macrophages were depleted with age together with InterFB and ParFB. Such macrophage dynamics can point to insufficient anti-inflammatory signals in the aged muscle after repeated rounds of damage and regeneration. Finally, both SMCs and pericytes decreased with age, which could lead to deterioration of the blood vessel wall, compromising blood flow to the muscle as has been previously suggested^84^.

To improve our understanding of ageing muscle microenvironment, we performed ageing DEGs analysis across microenvironment cell populations (Supplementary Table 3), paying particular attention to chemokines and cytokines. We also studied putative interactions mediated by these genes using the CellPhoneDB.org resource (Supplementary Table 6)^85^.

Strikingly, several cell populations in the ageing muscle either significantly up-regulated or had a tendency towards increased expression of *CCL2*. *CCL2* is the major pro-inflammatory and known monocyte/macrophage-attracting cytokine known to be activated in muscle injury (Fig. 5e)^86–88^. Indeed, CellPhoneDB analysis predicted that several microenvironment populations (FB, MuSC, ArtEC, SMC, pericytes) could attract monocytes, cDC2 and plasma cells to the ageing muscle via both *CCR2* and *CCR10* receptors (Fig. 5f). In addition to pan-microenvironment upregulation of *CCL2*, we also noted an increase in *CCL3*, *CCL4* and *CXCL8*, which was restricted to immune cells (Fig. 5e). For instance, *CCL3* and *CCL4*, that were found to be increasingly produced with age by CD14^+^ monocytes, macrophages and NK cells, were predicted to attract a whole range of immune cells including monocytes, macrophages, different types of DC, plasma and B cells as well as eosinophils and neutrophils (Fig. 5f, Extended Data Fig. 5e). Conversely, *CXCL8* was predicted to exclusively attract neutrophils (Fig. 5f). Taken together, this suggests a mechanism for increased immune infiltration into the muscle with age.

Finally, we noted an increase in expression of the pro-inflammatory cytokine *IL6* in several microenvironment cell types, coupled with a decrease in the anti-inflammatory *IL10* in immune cells (Fig. 5e)^89, 90^. Such an imbalance towards the inflammatory state could be both a cause and consequence of immune infiltration.

### Integrated human-mouse skeletal muscle ageing atlas at single-cell resolution

In order to compare and contrast age-related changes in human and mouse skeletal muscle, we integrated our in-house produced single-cell human and mouse skeletal muscle data with 6 previously published human^18, 21^ and mouse healthy, non-perturbed single-cell resources^19, 37, 39, 40^. The integrated dataset comprised 346,296 single-cells and contained samples from 33 human donors (19 to 84 years old) and 31 mice (1 month to 30 months) (Fig. 6a, Extended Data Fig. 6a-c, Supplementary Table 7).

**Fig. 6.**
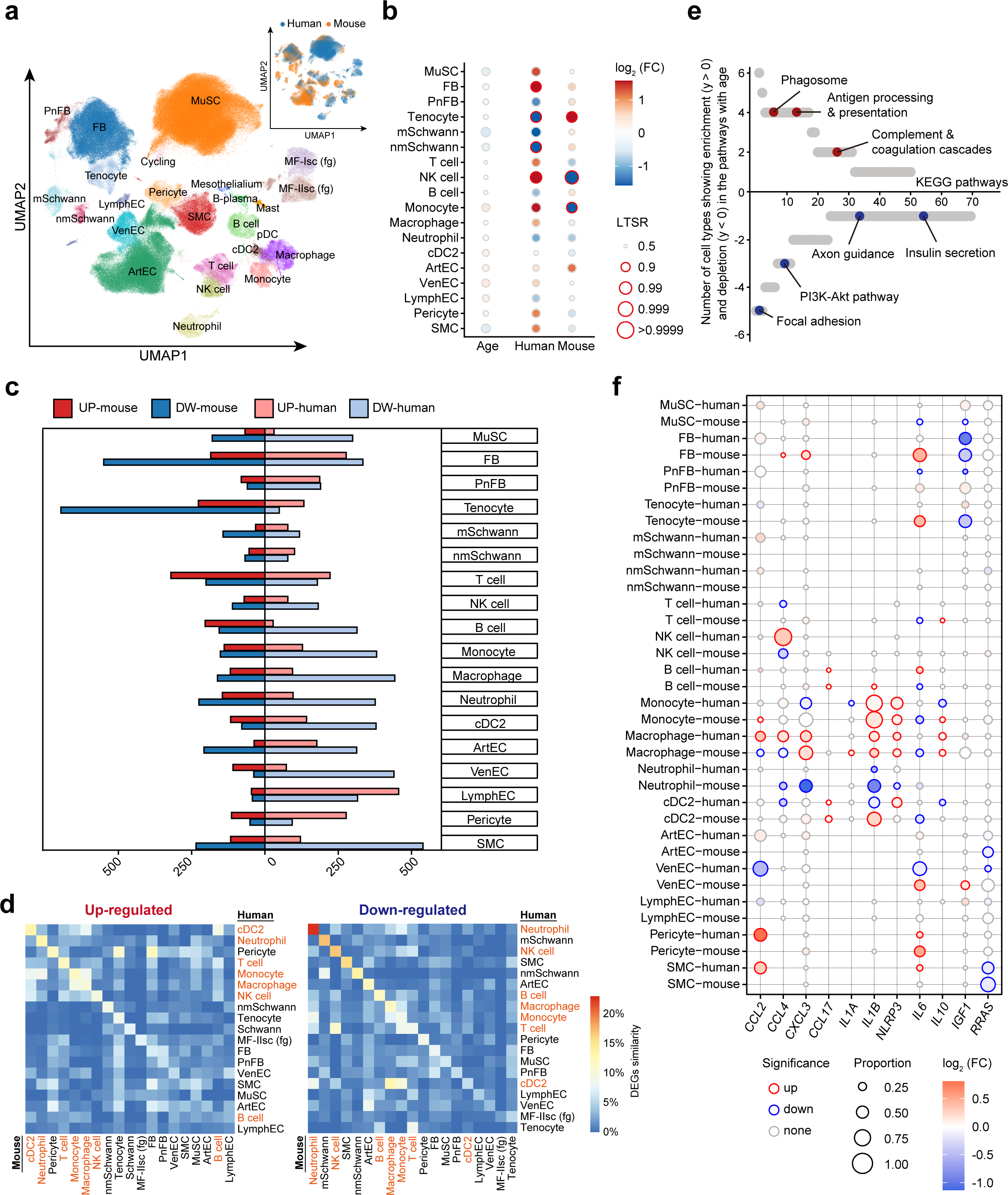
Integrated human-mouse skeletal muscle ageing atlas at single-cell resolution. **a**, UMAP plot showing main cell populations in the integrated human and mouse skeletal muscle dataset of 346,296 cells, including our muscle ageing atlas as well as 6 other publicly available resources. Insert shows UMAP plot coloured according to species. MF-Isc (fg) and MF-IIsc (fg), type I and type II myofiber fragment from scRNA-seq; MuSC, muscle stem cell; FB, fibroblast; EnFB, endoneurial fibroblast; PnFB, perineurial fibroblast; mSchwann, myelinating Schwann cell; nmSchwann, non-myelinating Schwann cell; B-plasma, plasma cell; ArtEC, arterial endothelial cell; VenEC, venous endothelial cell; LymphEC, lymphatic endothelial cell; SMC, smooth muscle cell. **b**, The log_2_ (FC) in the abundance of cell clusters with age (first column), and enrichment in human *vs.* mouse dataset (second and third columns), taking into account confounding factors. LTSR represents a significance measure indicating the local true sign rate. Significantly differentially abundant populations are highlighted with red edges (Methods). **c**, Barplot visualisation showing number of up- and down-regulated DEGs (x axis) in mouse and human across different cell types (y axis). **d**, Heatmap showing consistency in the DEGs between the same cell type in human and mouse for up- (on the left) and down-regulated genes (on the right). Consistency was calculated using Jaccard similarity index, immune cells are highlighted in red. **e**, Scatterplot illustrating number of cell types (y axis) which show simultaneous human and mouse age-related enrichment in the given KEGG pathway (x axis). Cell types showing enrichment in up-regulated genes are shown on the top (y > 0) *vs.* ones showing enrichment in down-regulated genes are displayed at the bottom (y < 0). KEGG pathways are ordered according to the number of cell types. **f**, Dot plot showing species-common and -specific ageing DEGs in human and mouse. Size of the dot represents the proportion of aged cells expressing the gene. Colour denotes log_2_ (FC) in aged *vs.* young upon gene expression. Significantly up-regulated genes are highlighted with red and down-regulated with blue edges.

To identify age-related changes in cell type abundance common to both human and mouse, we employed a linear mixed-effect Poisson regression model, while controlling for multiple covariates such as species effect, 10x chemistry of the library, muscle type and sex (see Methods). The common cross-species ageing effect was very weak, with a trend towards increase in monocytes, dendritic cells, ECs and decrease in MuSCs, Schwann cells and SMCs, though these changes were not statistically significant (LTSR < 0.9, see Methods) (Fig. 6b). At the same time, species had a much stronger effect on cell type abundance. MuSCs, fibroblasts, T cells, NK cells, monocytes, pericytes and SMCs tended to be more abundant in human, while tenocytes and ArtECs were more abundant in mouse (Fig. 6b). Having observed a weak agreement in the age-related changes in cell type abundance between species, we went on to compare cell type ageing DEGs between both species.

A systematic review of cell-type specific differential gene expression (see Methods) revealed on average a larger number of DEGs in human muscle compared to mouse (Fig. 6c, Supplementary Table 7). The majority of cell types showed more down-regulated genes with age than up-regulated ones (13 *vs.* 5 cell types for humans and 11 *vs.* 7 for mice), indicating that the predominant down-regulation of gene expression with age is conserved across species.

To further investigate the conserved ageing hallmarks at gene expression level, we determined the consistency of up-regulated and down-regulated genes between human and mouse using the Jaccard similarity index (see Methods). Consistency in ageing DEGs between human and mouse ranged between 1 and 19 % for different cell types (Fig. 6d). This is comparable to the 4.7% overlap we calculated using skeletal muscle ageing DEGs determined from bulk RNA-seq and reported by Zhuang et al. 2019^91^. Consistency of down-regulated ageing DEGs were larger than the up-regulated ones, further emphasising that down-regulation is a conserved ageing mechanism across species, as noted previously^92^. Furthermore, immune cells tended to have the highest cross-species overlap in the up-regulated genes as compared to other cell types (Fig. 6d, immune cells were highlighted in red). This likely indicates activation of gene expression programs which contribute to age-related inflammation. To investigate further, we performed KEGG pathway enrichment for ageing DEGs in human and mouse and plotted consistent ageing pathways between species in relation to the number of involved cell types (Fig. 6e). Immune-related pathways such as phagosome synthesis, antigen processing and presentation, complement cascade and coagulation were enriched in several cell types with age. This is in keeping with accumulated inflammation during muscle ageing^93, 94^. Our data also revealed sets of pathways maintaining healthy muscle functions to be depleted in multiple cell types with age, including the PI3K-Akt pathway, focal adhesion and axon guidance (Fig. 6e, Supplementary Table 7), further indicating a general decline in muscle physiology.

Through analysing the DEGs in each cell type individually, we revealed a subset of genes potentially driving muscle ageing in both the human and mouse (Fig. 6f). Of note, the increase in the immune-attracting cytokine *CCL2 and CCL4* we reported in the human skeletal muscle was not replicated in the mouse single-cell data (Fig. 6f). Pan-cell-type increase of *CCL2,* in particular, may be a specific feature of human muscle ageing. At the same time, other pro-inflammatory molecules such as chemokines *CXCL3*, *CCL17* and interleukin *IL1B* together with inflammasome *NLRP3* showed increase in both species’ monocytes, macrophages, cDC2 and B cells (Fig. 6f, Supplementary Table 7), pointing to an increase in inflammation with age.

While we observed proinflammatory *IL6* tended to increase with age in pericytes and smooth muscle cells of both species, it has also shown a lot of decrease in mouse as opposed to human (MuSC, T cells, B cells, monocytes, macrophages and cDC2, for mouse and VenEC, PnFB for human) (Fig. 6f). Anti-inflammatory *IL10* was increased in both human and mouse macrophages (Fig. 6f), but decreased in human monocytes and cDC2 whilst increasing in mouse monocytes. Finally, growth and angiogenesis associated with *IGF1* and *RRAS* genes were mostly down-regulated in stromal and blood vessel cells, respectively, both in humans and mice.

Taken together, this suggests an increase in inflammation coupled with decreased growth signalling is the common feature in skeletal muscle ageing of both species. However, it may be orchestrated by different cytokines and chemokines across different cell populations.

## Discussion

While single-cell genomics studies have provided many insights into ageing in rodent tissues, studies in human tissue have been limited. The small number of human studies have yielded interesting insights on, for instance, ageing of pancreas^95^, skin^96^ and retina^97^. The bottleneck for human studies is the limited access to healthy human tissue across the lifespan. Additionally, in the specific case of skeletal muscle, comprehensive single-cell studies have the technical challenge that the syncytial myofiber is impossible to profile with droplet single-cell technologies due to its size, unlike MuSCs and resident muscle environment cells. Here, we combined single-cell and single-nucleus RNA sequencing to build the first human skeletal muscle ageing atlas that includes both MuSCs, microenvironment cells and myofiber nuclei. In this atlas, we annotated 39 major cell types, providing more fine-grained insights than was previously available^18, 19, 23, 24, 37^ .

From this detailed analysis, we identified novel myonuclei populations, and hypothesised that their presence and change in abundance may be responsible for the differential ageing of slow-twitch and fast-twitch myofibers. Importantly, we unveiled two compensatory mechanisms for fast-twitch or glycolytic muscle atrophy, one mediated by the glycolytic-shifting of the oxidative nuclei, and another one by newly generated MYH8^+^ myocytes from MuSCs. We also revealed novel specialised NMJ accessory myonuclei which can facilitate rebuilding of the dysfunctional NMJ.

In the myofiber microenvironment, we found several cell types which displayed pro-inflammatory changes, expressing pro-inflammatory chemokines such as *CCL2, CCL3, CCL4*. Their interactions with immune cells may be responsible for the infiltration of immune cells and chronic inflammation in the aged muscle. Finally, our human-mouse integrated ageing atlas helps to reveal human-specific cellular and regulatory mechanisms of muscle ageing.

It is well-known that fast-twitch myofiber is more susceptible to atrophy in ageing compared with slow-twitch ones. Several mechanisms have been proposed as explanations for this phenomenon^68, 98^. Through snRNA-seq we identified several types of nuclei which are differentially distributed in slow-twitch and fast-twitch myofibers, and can be related to fast-twitch myofiber degeneration (TNFRSF12A^+^) and slow-twitch myofiber regeneration (OTUD1^+^ and FAM189A2^+^), respectively. Therefore, we hypothesise that fast-twitch myofiber atrophy in ageing is intrinsic to myofiber and controlled by the subpopulations of the nuclei. Our results also demonstrate the plasticity of gene-expression profiles in single nuclei in syncytial cells, which can adapt under physiologic or pathologic stress. This may contribute to the resilience of skeletal muscle to ageing, metabolic change, and exercise.

Additionally, loss of muscle mass and function in ageing could be due to the dysfunction of innervation by the peripheral nervous system, caused either by degeneration of the myofiber apparatus linking to motor neurons, or impairments of the neurons themselves. In our work, we could not detect changes in the abundance of NMJ nuclei, which are responsible for the formation of NMJ apparatus, over ageing. However, we did find a significant decrease of both myelinating and non-myelinating Schwann cells, which normally served to protect and support neurons. Therefore, we suggest that the degeneration of NMJ in muscle ageing can at least partially be attributed to the loss of neural function as a consequence of glial cell loss. Importantly, we observed an increase in novel NMJ accessory myonuclei in aged myofibers, especially of the slow-twitch type, which may help re-innervate the NMJ. Although this physiological compensatory mechanism seems to ultimately fail in preventing muscle atrophy, it might provide a future avenue for medical intervention to improve muscle function.

Overall, our integrated genomics, imaging and computational analyses provide a global overview of the biology of muscle ageing and can serve as a valuable resource for future human ageing studies.

## Materials and Methods

### Tissue acquisition

Human intercostal muscle samples (inner part between 2^nd^ and 3^rd^ ribs) for single-cell and nuclei processing were collected with consent from deceased transplant organ donors by the Cambridge Biorepository for Translational Medicine, Cambridge, UK (CBTM), immediately placed in HypoThermosol FRS preservation solution and shipped to Sanger Institute for processing. Ethical approval was granted by the Research Ethics Committee East of England - Cambridge South (REC Ref 15/EE/0152) and informed consent was obtained from the donor families. Three 19 months old and five 3 months old male mice of C57BL/6JRj strain were obtained from Janvier labs, France and housed at the Sanger Institute under Establishment licence number X3A0ED725 provided by the Home Office. They were used to dissect hindlimb muscles for the following single-cell and single-nuclei isolation.

For experimental validations, adult human intercostal muscle biopsies were collected from Sun Yat-sen Memorial Hospital (Guangzhou, China) under approval of the Research Ethics Committee of Sun Yat-sen University (REC 2018-048). For isolation of human primary myoblasts, embryonic hindlimb muscles were collected from medical abortions at Guangzhou Women and Children’s Medical Center (Guangzhou, China) with ethical approval license (REC 2019-075) granted by the Research Ethics Committee of Sun Yat-sen University. Informed consent for intercostal and fetal muscle tissue collection and research was obtained from every patient. Collected intercostal muscles were either freshly processed for immunohistochemistry assays or used for FACS-sorting to isolate MuSC subpopulations. Fetal hindlimb muscles were enzymatically digested to purify myoblasts for *in vitro* culturing.

### Single-cell sample processing

Skeletal muscle tissue was processed according to the following protocols for single-cell and single-nucleus isolation from skeletal muscle deposited at protocols.io^99, 100^. Briefly, muscle tissue was minced, digested in the solution of Collagenase II (Worthington-Biochem, LS004176) and Dispase (Gibco, 17105041) and, following that, lysate was centrifuged in the gradient of Percoll to recover a fraction with single-cells. For single-nucleus isolation, muscle tissue was grounded using a dounce homogenizer, lysed in the nuclei lysis buffer and, finally, Percoll gradient was used to separate intact nuclei and cell debris. For the scRNA-seq experiments, either 8000 live cells or 8000 intact nuclei were loaded per sample into Chromium Controller (10x Genomics) and Single Cell 3’ v2 or v3 Reagent Kit was used to create GEM, perform cDNA synthesis and generate sequencing libraries. The libraries were sequenced on an Illumina HiSeq 4000 or Novaseq 6000 with the following parameters: Read1: 26 cycles, i7: 8 cycles, i5: 0 cycles; Read2: 98 cycles to generate 75-bp paired-end reads.

### Transcriptome mapping

3’ 10x Genomics skeletal muscle droplet sequencing data for human and mouse cells were aligned and quantified using the Cell Ranger Single-Cell Software Suite version 3.1.0 and GRCh38 human or mm10 reference genome (official Cell Ranger reference, version 1.2.0). Corresponding single-nucleus data for mouse and human were aligned and quantified with Cell Ranger made pre-mRNA reference genomes. Spliced and unspliced counts were derived from Starsolo analysis^101^ using GRCh38-3.0.0 human genome and parameters matching Cell Ranger 3.x.x results.

### Cell quality control and filtering

Cellbender^102^ 0.2.0 was used to remove ambient RNA contamination from both single-cell and single-nucleus data with the following parameters: n_epochs = 150 and learning rate 0.0001 (for some samples these were adjusted to 250 epochs and 0.00005 learning rate).

Scrublet^103^ was used to identify potential doublets in each sample and after that, cells with scrublet score > 0.4 were filtered out as doublets. Next, additional filtering was performed to discard potential empty droplets and cells with high percentage of ambient RNA using the following minimum thresholds: 1000 UMI and 500 genes for cells and 700 UMI and 500 genes for nuclei. Cells with higher than 20% and nuclei with more than 5% mitochondrial genes expressed were removed as potential low-quality cells.

### Data processing and batch alignment

Scanpy python package (version 1.7.2)^104^ was used to load the cell by gene count matrix and perform processing according to standard pipeline with modifications. First, we normalised raw gene read counts by sequencing depth using *scanpy.pp.normalize_per_cell* followed by ln(x) + transformation, done using *scanpy.pp.logp*. Then, we selected 3000 or 10000 highly variable genes using *scanpy.pp.highly_variable_genes (flavor=’seurat_v3’),* which mimics the procedure implemented in Seurat v3. Following that, we performed dimensionality reduction and batch correction on the data using scVI model (parameters: n_layers = 2, n_latent = 30) from the package scvi-tools^27^. We used 10x library as a batch and specified data modality (cells *vs.* nuclei), 10x chemistry, donor ID, sex as additional categorical and percent_mito as numerical covariates to correct for. Next, we calculated the neighbourhood graph using *scanpy.pp.neighbours* with k = 15 and used it to perform cell clustering with the Leiden algorithm (resolution = 1).

Marker genes were identified using different approaches. In most cases, *t*-test was applied to identify DEGs in the given cluster as compared to the rest using *sc.tl.rank_gene_groups* (method = ‘t-test_overestim_var’, corr_method = ‘benjamini-hochberg’), obtained *P* values were corrected using benjamini-hochberg method and top 100 genes were considered. Alternatively, *scvi.model.SCVI.differential_expression* function was used to calculate differentially expressed genes (DEGs) between a particular cluster and the reference using Bayesian approach. This was used to better call markers for specialised nuclei populations such as I-FAM, I-OTU, II-FAM, II-OTU and II-TNF. Specifically, DEGs were called by comparing every specialised population to the conventional type I or type II myofiber cluster, respectively. Later, DEGs were further prefiltered to have log_2_ (Fold change, FC) above one, to be expressed in at least 10% of the cells and to have Bayes factor above two. Finally, gene set over-representation analysis was performed on the marker genes either using *scanpy.queries.enrich*, a wrapper for gprofiler^105^, or Metascape web-interface^106^ .

Trajectory analysis aimed to uncover intermediate stages between MuSC and myofiber was performed using the Monocle2 R package (v2.9.0)^107^. Dataset for trajectory analysis was limited to MuSCs and myofiber cells (coming from scRNA-seq) and the known genes, important for muscle differentiation: *PAX*, *MYF5*, *PDGFRA*, *MYOG*, *TPM1*, *MYH2*, *MYH3*, *NCAM1*, *TNNT1*, *TNNT2*, *TNNT3*, *TNNC1*, *CDK2*, *CCND1*, *CCNA1*, *ID1*.

### Integration of publicly available datasets

With the aim to compare quiescent and activated human and mouse MuSC subtypes, we have downloaded and integrated mouse MuSC from the following resources: GSE110878, GSE143476, GSE134540, GSE138707 and GSE149590.

With the aim to create an integrated human-mouse skeletal muscle dataset we have identified and downloaded all healthy unchallenged human and mouse skeletal mouse datasets, which were publicly available. The count matrices for human skeletal muscle scRNA-seq samples were obtained from Gene Expression Omnibus (GEO) GSE143704 and DRYAD (https://doi.org/10.7272/Q65X273X), while corresponding mouse samples were obtained from the following repositories: GSE110878, GSE138707, GSE134540, GSE143476, GSE149590, GSE142480. All human and mouse samples underwent Scrublet doublet detection and QC filtering as described previously. Next, human and mouse datasets were concatenated into one object, retaining only homologous genes. After standard pre-processing, the scVI model was used to produce an integrated embedding for the data, while using SampleID as a batch and species, 10x_chemistry, muscle type, Sex as categorical covariates.

#### Ageing cell type composition analysis

With the aim to identify the age effect on cell composition we have used two different approaches depending on the size of the dataset and distinctness of the clusters.

1. Mixed-effect Poisson regression model was used to assess the effect of age on major human and mouse cell type compositions, while controlling for confounding covariates.

Specifically, the effect of age on cell type specific counts was modelled using the Poisson linear mixed-effect model accounting for the possible biological and technical covariates using *glmer* function from the “lme4” package on R. We provided all of the factors as mixed terms (for instance, 1|X) as these allow estimation of the coefficients despite the collinearity of covariates. The effect of age and most of the covariates were estimated as an interaction term with the cell type. The log-transformed fold change for every covariate was calculated relatively to the grand mean and adjusted so its value is 0 when there is no effect. Local true sign rate (LTSR) was used to estimate statistical significance, which denotes probability that the estimated direction of the effect is true

(see details on its calculation here^108^ Cell type composition analysis). LTSR ranges from 0 to 1, where the higher value denotes higher probability, we used LTSR > 0.9 as a cutoff to call significant age effect on cell type compositions.

- The model fitted for major human cell types is (Fig. 1c): *Ncs ∼ Age + (1|Celltype) + (1|Sample) + (1|10x) + (1|batch) + (1|Sex) + (Age-1|Celltype) + (1|10x::Celltype) + (1|batch::Celltype) + (1|Sex::Celltype)+ (1|Sample::Celltype),* where *Ncs* denotes the cell count of cell type c in sample n, *Age* denotes age in years, scaled and centered, *Sample* denotes 10x library ID, *batch* denotes cells or nuclei.
- The model fitted for combined human-mouse dataset (Fig. 6b): *Ncs ∼ (1|Age_group) + (1|Celltype) + (1|Sample) + (1|10x) + (1|Species) + (1|Sex) + (1|Age_group::Celltype) + (1|10x::Celltype) + (1|Species::Celltype) + (1|Sex::Celltype) + (1|Sample::Celltype),* where *Ncs* denotes the cell count of cell type c in sample n, *Age_group* denotes binary age (young, old), *Sample* denotes 10x library ID and *Species* denotes human or mouse.

2. MiloR (v1.2.0)^41^ R framework was used to detect ageing changes in the states within one or several cell types (for MuSC, myofiber, immune, fibroblast and vasculature cell types).

We preferred the Milo approach for smaller scale datasets as it provides more resolution for the change and is not limited by the resolution of clusters/cell types. Specifically, we first constructed a KNN-graph of cells (k was adjusted depending on cell type, d = 30) using embedding obtained after application of the scVI model on a particular cell type(s). Next, we sampled a representative group of cell neighbourhoods across KNN-graph and obtained a count matrix with neighbourhoods in rows and sample ids in columns. Following that, we applied a negative binomial model linear regression model to assess the effect of Age on the number of cells in each neighbourhood accounting for the 10x chemistry and Sex effect. The significance was controlled for multiple testing using weighted BH correction^41^. We later assigned neighbourhoods to cell type labels based on the majority voting of the cells comprising that neighbourhood, where the most abundant label was present in more than 70% of cells. Otherwise, the neighbourhood was assigned a “Mixed” label.

### Ageing differential gene expression analysis

We performed ageing differential gene expression analysis twice to identify age-associated genes changing in every major cell type as well as in every subtype. With that aim we employed the linear mixed model as proposed^109^, which allowed us to account for various technical (10x chemistry, data modality: scRNA-seq or snRNA-seq) and biological covariates (Donor, Sex) to disentangle the true effect of age. After fitting the model, a Bayes factor of each factor was calculated for every gene as described in the previous report^108^. Later Bayes factors were used to compute posterior probability and significance measure, LTSR (see section 1.3 of the supplementary note^109^ for more details) for the influence of the factor on every gene. The same model was also applied to test for age-associated DEGs separately within each species (human, mouse) while using muscle type, donor/mouse sex as biological covariates and 10x chemistry and data modality (scRNA-seq or snRNA-seq) as technical covariates.

To compare ageing DEGs between species, we calculated the Jaccard index, which denotes the ratio between the size of the intersection of two sets and their union. Next, we used the “clusterProfiler” R package to identify which KEGG pathways are enriched among ageing DEGs in every species. We used genes that have log_2_ (FC) above 0 or below 0, LTSR greater than 0.9 and were expressed at least in 5% of cells in the relevant cell type to perform enrichment analysis. After selecting significantly enriched pathways, we scored them based on the number of cell types that showed simultaneous human and mouse age-related enrichment (for log_2_ (FC) > 0) and depletion (for log_2_ (FC) < 0) in them.

### Cell-cell communication analysis

We used the CellPhoneDB algorithm (v3)^85^ and database to obtain the list of all possible cell type pairs and receptors which can interact through the following ligands *CCL2*, *CCL3*, *CCL4* and *CXCL8*. Normalised count data and broad cell type assignment were used as input. Only receptors and ligands expressed in more than 5% of the cells in the specific cluster were considered to indicate relevant interactions. Next, we used the “igraph” R package to visualise all possible emitter cell types, producing the aforementioned ligands and all possible receptor cell types, expressing the receptor. We also indicated if the expression of ligand or receptor has changed with age (according to ageing DEGs analysis).

### FACS-based cell sorting of MuSC

Freshly obtained intercostal muscles were collected in sterile 1 × PBS. The superficial connective tissue and blood contaminants were carefully removed using forceps and scissors under the stereo microscope. Tissue was then mechanically dissociated with fine scissors in 10 mL (per gram of tissue) enzyme solution mixed with 2.5 U/mL Dispase II (Roche, #4942078001) and 1 mg/mL Collagenase B (Roche, #11088815001) supplemented with 5 mM MgCl_2_ and 2% penicillin–streptavidin (Gibco, #15140122). Minced tissue was digested with enzyme solution at 37°C for 60-90 min with gentle shaking, filtered sequentially through 100 µm (Falcon, #352360) and 40 µm (Falcon, #352340) cell strainers to get the single cell suspensions. Cell suspensions were adjusted to 2-7.5 × 10^6^ cells/mL with FACS buffer (2% FBS diluted in 1 × PBS) and incubated with the following fluorophore-conjugated antibodies: anti-human CD31-PE monoclonal antibody (eBioscience, #12-0319-42, 1:200 dilution) for negatively separating endothelial cells, anti-human CD82-PE/Cyanine 7 monoclonal antibody (BioLegend, #342109, 1:500 dilution) and anti-human CD56-PE/Cyanine 7 monoclonal antibody (eBioscience, # 25-0567-42, 1:200 dilution) to enrich human MuSCs^42, 43^, and anti-human CD54-APC monoclonal antibody (eBioscience, #17-0549-41, 1:300 dilution) for sorting ICA^+^ MuSCs. Cells were incubated on ice for 30 min, while being protected from light. After washing with FACS buffer, cells were sorted and analysed with BD Influx^TM^ Cell Sorter. Exported raw data were processed with FlowJo (10.4) to analyse cell populations.

### RNA isolation for sorted cells and quantitative PCR

FACS-sorted ICA^+^ MuSC and ICA^-^ MuSC either from young or aged muscle biopsies were collected directly into 1 mL of TRIzol reagent (Invitrogen, #15596026) for isolation of total RNA. After quality control and quantification by NanoDrop^TM^ One (Thermo Scientific), 100 ng of RNA was reverse transcribed into cDNA using PrimerScript^TM^ RT Master Mix (TaKaRa, #RR036A). Next, 100 ng cDNA from each sample was subjected to quantitative real-time PCR (qPCR) following the manufacturer’s instructions of PerfectStart Green qPCR SuperMix (TransGen Biotech, #1) on LightCycle480 Instrument II (Roche). Relative mRNA expression was normalised to *RPLP0* using standard statistics method of 2^-ΔΔCT^.

### Immunohistochemistry

Freshly collected intercostal muscles were trimmed with forceps to remove the superficial connective tissue, embedded into Tissue-TEK^®^ OCT compound (PST) and frozen in isopentane cooled with liquid nitrogen. For immunohistochemistry, 10 µm fresh-frozen cryosections were collected from the OCT-embedded muscle tissue and fixed with 4% PFA. Sections were then incubated in citrate buffer (pH 6.0) at pressure cooker to perform heat activated antigen retrieval. After washing with 0.1% Triton X-100 diluted in PBS (0.1% PBST), cryosections were blocked with 10% affinipure Fab goat anti-mouse IgG (Jackson Immunoresearch, #115-007-003) in 0.1% PBST for 60 min and 5% normal goat serum (Jackson Immunoresearch, #005-000-121) in 0.1% PBST containing 2% BSA for 30 min, respectively. For immunostaining, the sections were incubated with the following mixed primary antibodies overnight at 4°C: ant-MYH7 (Developmental Studies Hybridoma Bank, DSHB, #BA-F8, 1:14 dilution), anti-MYH2 (DSHB, #SC-71, 1:20 dilution), anti-MYH1 (DSHB, #6H1, 1:6 dilution) or anti-MYH8 (DSHB, #N3.36, 1:9 dilution). After washing with 0.1% PBST, sections were incubated with mixed secondary antibodies at room temperature for 1 hour and later with DAPI for 10 min both at room temperature. Mix of secondary antibodies included goat anti-mouse IgG1 (Alexa Flour 488, Invitrogen, #A-21121, 1:400 dilution), goat anti-mouse IgG2b (Alexa Flour 647, Invitrogen, #A-21242, 1:400 dilution) and goat anti-mouse IgM (Alexa Flour 555, Invitrogen, #A-21426, 1:400 dilution). After staining, sections were covered with fluorescence-saving mounting medium (Millipore, #345789) and imaged with DMi8 inverted microscope (Leica Microsystems). For presentation, the MYH7^+^ colour channel was adjusted from purple to blue and MYH8^+^ colour channel from grey to red to better show merged images.

### Image analyses

For unbiased myofiber subtyping and age-associated comparisons (Fig. 4b-d, Extended Data Fig. 4b-e, Supplementary Table 5), we developed an automated myofiber segmentation and image analysis workflow. Briefly, for each multi-channel image exported from the microscope, all channels targeting non-nuclei channels were first max-projected and then sequentially processed through gaussian blurring, gamma adjustment and USM^110^. After that, myofiber segmentation was performed using a deep-learning based object segmentation algorithm, Cellpose^111^. In order to classify myofibers into different fast and slow subtypes, we have manually trained an object classifier using the object classification workflow in ilastik^112^. This classifier also returns metrics that describe myofibers’ properties which were later exported for downstream analysis. To ensure trivial myofiber quantification, all myofibers located at the image border are discarded. Nuclei surrounding all myofibers were also detected with Cellpose using a different set of parameters in both preprocessing and segmentation steps (see Extended Data Fig. 4a for details).

For automatic analysis of MYH8^+^ myofiber area done on teased muscle pieces (Fig. 4j, Supplementary Table 5), we used Fiji to quantify size in pixels of both the whole muscle pieces and MYH8^+^ myofiber areas. Multiple images from each sample were stitched with Fiji stitching tool to obtain the whole muscle image. Later, area in pixels occupied by the tissue was quantified as “whole muscle area”, while “MYH8^+^ area” was quantified as an area in pixels above the default threshold within the green channel (MYH8^+^ channel). Finally, we detected and quantified centralised nuclei manually (Figure 4l, Supplementary Table 5), since automatic approaches were not very accurate.

### Immunofluorescence on teased human skeletal muscles

Fresh muscle samples were trimmed to get rid of the superficial connective tissue under stereomicroscope and then immediately fixed with 4% paraformaldehyde (PFA) at room temperature for 8 min. Freshly fixed muscles were gently teased with forceps into thinner pieces in PBS and fixed with 4% PFA for another 10 min. After washing with 0.5% Triton X-100 diluted in PBS (0.5% PBST), muscle pieces were then transferred into PBS and incubated at 55°C for 30 min to denature the extracellular collagen fibrils. Next, muscles were blocked with 5% normal goat serum (Jackson Immunoresearch, #005-000-121) diluted in 0.5% PBST containing 3% BSA for 60 min. Immunofluorescence was performed with the following primary antibodies: anti-NEFH (Cell Signaling Technology, #2836S, 1:400 dilution), anti-S100B (Abcam, #ab52642, 1:200 dilution), anti-SORBS2 (Proteintech, 24643-1-AP, 1:200 dilution), and anti-MYH8 (DSHB, #N3.36, 1:9 dilution). For AChRs staining, Cy3 conjugated α-Bungarotoxin (α-BTX, BosunLife, #00018) at a final concentration of 2.5 µg/mL (performed under the Biosafety Cabinet) was mixed with primary antibodies and applied overnight at 4°C. Muscle pieces were then washed with 0.5% PBST and incubated with fluorescein-conjugated secondary antibodies including: goat anti-mouse IgG H+L (Alexa Flour 488, Invitrogen, #A-11029, 1:400 dilution), goat anti-rabbit IgG H+L (Alexa Flour 488, Invitrogen, #A-11008, 1:400 dilution), goat anti-mouse IgM (Alexa Flour 555, Invitrogen, #A-21426, 1:400 dilution). Nuclei were stained with DAPI (Invitrogen) and muscle bundles were then covered with fluorescence-saving aqueous mounting medium (Millipore, #345789). Images were taken with Nikon C2 Confocal Microscope (Nikon Eclipse Ni-E) and DMi8 inverted microscope (Leica Microsystems). Due to NMJs position only in one specific part of the myofiber (in the middle) of myofiber, it is often hard to capture them in the myofibers from surgical biopsies. We could only detect NMJ structures in one young and two aged muscle biopsies among over thirty biopsies surveyed. Hence, our statistics are based on those samples (Fig. 3h, i).

### Dissociation of human primary myoblast and cell culture

Human embryonic hindlimb muscles were collected in sterile PBS and mechanically dissociated using fine scissors in 10 mL (per gram of tissue) of enzyme solution containing 2.5 U/mL Dispase II (Roche, #4942078001) and 1 mg/mL Collagenase B (Roche, #11088815001) supplemented with 5 mM MgCl_2_ and 2% penicillin–streptavidin (Gibco, #15140122). Tissue was incubated with enzyme solution for 30 min at 37°C with gentle shaking. Digestion was stopped with 10% FBS plus 2 mM EDTA diluted in sterile PBS and tissue suspension was sequentially filtered through 100 µm (Falcon, #352360) and 40 µm (Falcon, #352340) cell strainers to get the single cell suspension. After centrifugation and repeated washing, cells were pre-plated in a 10 cm Gelatin-coated cell culture dish for 40 min. This procedure allows the majority of the fibroblasts to adhere to the dish ahead of myoblasts. After pre-plating, the cell supernatant was then gently transferred to a new Gelatin-coated cell culture dish to enrich primary myoblasts under the conditions of 5% CO_2_ and 37°C.

Purified myoblasts were grown in DMEM/F-12 cell culture medium containing 20% FBS, 10 ng/mL human basic fibroblast growth factor (PeproTech, AF-100-18B-500) and 1% penicillin-streptavidin. For myogenic induction, cells at 80% confluence were moved to the differentiation DMEM/F-12 medium containing 2% house serum and 1% penicillin-streptavidin. Myoblasts were differentiated to myotubes within 2-3 days.

### siRNA-mediated gene knockdown assay

siRNA oligonucleotides specifically targeting *SORBS2* transcripts were designed using the online toolkit (https://rnaidesigner.thermofisher.com/rnaiexpress/). Synthesised siRNAs were diluted with sterile RNase-free ddH_2_O to reach 20 mM stock concentration and stored at -20°C. For siRNA transfection, 5 µL of 20 mM siRNAs and 3 µL of X-tremeGENE HP DNA Transfection Reagent (Roche, #6366546001) were diluted in 200 µL Opti-MEM (Gibco, #11058021) and incubated for 15 min to allow transfection complex formation. Later the mix was slowly added to the cell culture medium in each well of 6-well plate to reach a final siRNA concentration of 50 nM. 48 hours after transfection, cells were assigned either to qPCR to detect knockdown efficiency or to immunofluorescence to evaluate effects on AChRs aggregation formation.

### In vitro induction of AChRs aggregation

Primary human myoblasts were seeded in 6-well cell culture plates for differentiation and AChRs aggregation induction. Before seeding cells, plates were pre-coated with 10 µg/mL natural mouse laminin (Gibco, #23017015) diluted in DMEM/F-12 at 37°C for at least 4 hours, which is essential for postsynaptic apparatus formation *in vitro*^113, 114^. Upon reaching 80% confluence, cells grown in laminin-coated plates were switched for differentiation. On day 2 of myogenic differentiation, cells were first incubated with 200 µL of 10 µg/mL laminin for 20 min and later supplemented with 2 mL of differentiation medium for continuous culturing in order to induce formation of mature AChRs clusters. Once myoblasts got differentiated into myotubes on day 3, siRNAs targeting *SORBS2* transcripts were transfected into the myotubes. 48 hours post transfection, myotubes were fixed with 4% PFA, washed with 0.5% PBST and stained with 2 µg/mL α-BTX followed by DAPI. Topological AChRs aggregates were visualised using DMi8 inverted microscope (Leica Microsystems). Quantifications of AChRs clusters were carried out with help of Fiji software.

### Statistics

In this study, all of the bioinformatic analyses related to sc-/snRNA-seq data processing were described in detail in the Methods and corresponding figure legends. For all of the plots in the study, number of biological repeats, detailed statistical methods and significance were stated in the Methods and corresponding figure legends.

## Data availability

The processed data objects generated within this study are available for browsing and inspection by the editors and reviewers at the following website https://www.muscleageingcellatlas.org. Raw sequencing data for the newly generated libraries will be deposited to Array Express and accession codes will be available before publication. The publicly available human skeletal muscle scRNA-seq data were downloaded from the GSE143704 and DRYAD (https://doi.org/10.7272/Q65X273X) repositories while mouse scRNA-seq samples were obtained from GSE110878, GSE138707, GSE134540, GSE143476, GSE149590, GSE142480 repositories.

## Code availability

All code scripts and notebooks used in the study are available upon request and will be deposited at GitHub repository before publication.

## Acknowledgments

We are grateful to the donors, their families and the Cambridge Biorepository for Translational Medicine for the gift of tissue. We acknowledge Chenqu Suo for assistance with muscle single-cell processing, Cecilia Domínguez Conde and Kelvin Tuong as well all of the members of the immune subgroup of Teichlab for advice on immune cells, Aidan Maartens for proofreading this manuscript, Martin Prete for creating the website, Christina Usher and BioRender.com for graphical illustrations and members of sequencing pipelines at Sanger for sequencing samples and aligning the data. We are also deeply grateful to all members of Teichlab and Zhang groups for interesting discussions and their thoughtful comments.

The research in H.Z. laboratory is supported by the National Key R&D Program (grant number: 2019YFA0801703), National Natural Science Foundation of China (grant number: 31871370 and 32000840) and Advanced Medical Technology Center of Sun Yat-sen University (K0507007). We also acknowledge funding provided by Wellcome Human Cell Atlas Strategic Science Support (WT211276/Z/18/Z). S.A.T.’s research is funded by Wellcome (WT206194) and the ERC Consolidator Grant ThDEFINE.

## Author Contributions

H.Z. and S.A.T. conceived the study and supervised the work. K.T.M., K.S.P. acquired organ donor tissue and Z.S., A.P.X collected biopsies from human patients. Y.W., H.Z., M.D., V.R.K, C.T., M.P. preserved collected tissue. L.B., H.Z., E.S.F., E.P., V.R.K performed single-cell and nuclei processing of the tissue. Y.W. devised *in vitro* and *in vivo* validation strategies, established experimental workflows, performed wet-lab experiments and data interpretation. X.C. helped Y.W. with tissue pre-processing, Q.G. assisted Y.W. with immunohistochemistry and X.L. provided FACS instrumental support.

V.R.K., H.Z., S.A.T. coordinated data analysis across two sites and shaped study direction with input from Y.W. and T.Liu. V.R.K, V.K., N.K., N.H. performed statistical method development for the project. Data processing: K.P., V.R.K.; Global analysis: V.R.K, T.Liu.; MuSC analysis: T.Liu, Y.W., V.R.K.; Myofiber analysis: V.R.K, Y.W., T.Liu, Immune analysis: V.R.K., T.Liu, Vascular analysis: V.R.K, L.Y., T.Liu, Fibroblast analysis: V.R.K, T.Liu., cross-species integration and mouse data analysis: T.Liu, V.R.K, cross-species ageing analysis: V.R.K. Myofiber segmentation and feature extraction: T.L. and O.A.B. ; manual image analysis and myofiber type quantification: Y.W. Interpretation of the results: V.R.K, Y.W., H.Z., S.A.T, T.Liu with help of J.S.L and K.B.M. Manuscript writing: V.R.K., Y.W., H.Z., T.Liu, J.S.L., S.A.T, K.B.M.

## Competing interests

In the past 3 years, Sarah A. Teichmann has consulted for Genentech and Roche and sits on Scientific Advisory Boards for Qiagen, Foresite Labs, Biogen, and GlaxoSmithKline and is a co-founder and equity holder of Transition Bio.

## Extended Data Figure Legends

**Extended Data Fig. 1.**
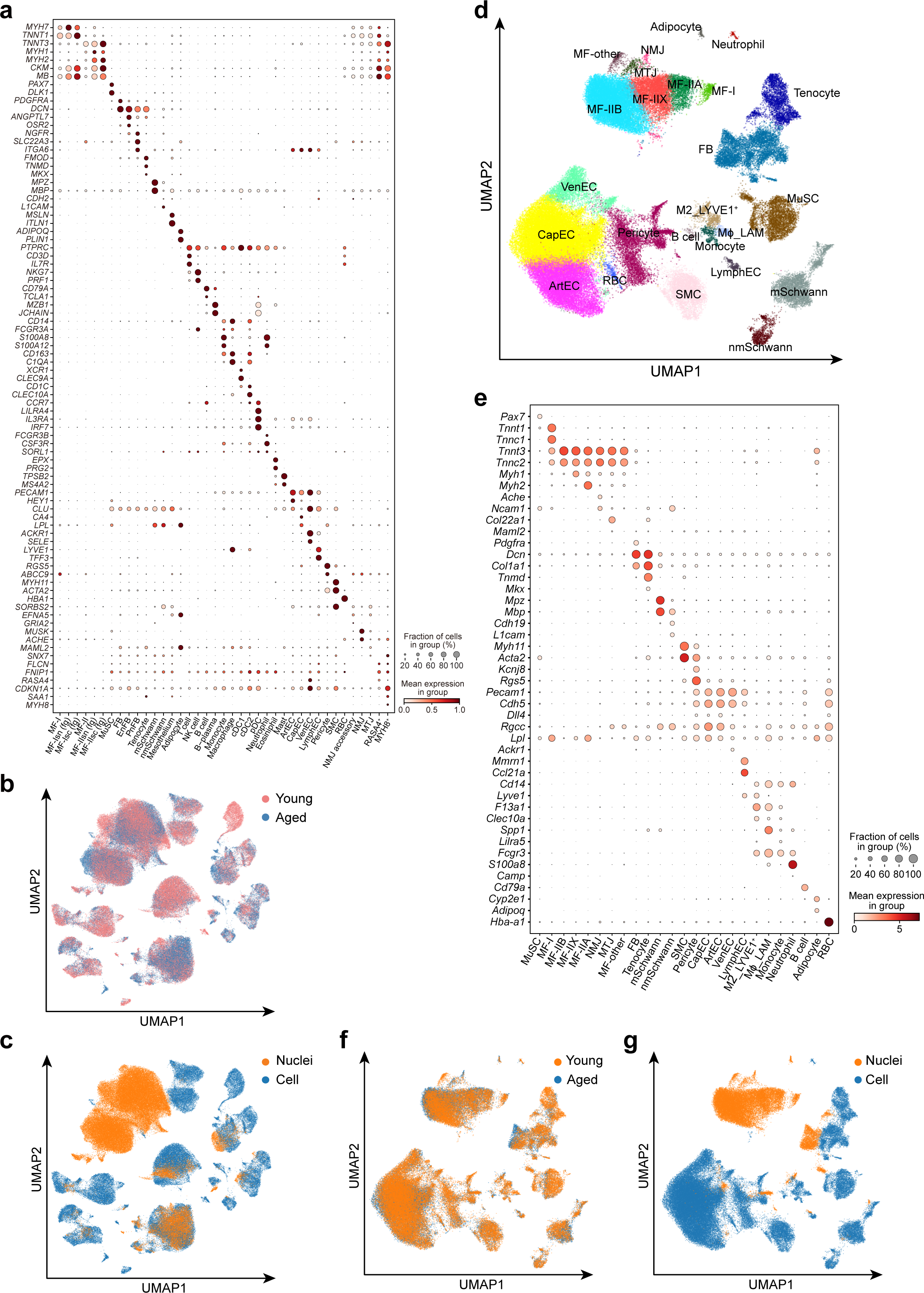
A single-cell and single-nucleus transcriptomic atlas of human skeletal muscle ageing. **a**, Dot plot showing marker genes for major cell types in human skeletal muscle ageing atlas. Size of the dot represents the proportion of cells expressing a gene. Colour denotes scaled expression level. **b**, **c**, UMAP visualisation of human ageing cell atlas coloured according to age (young *vs.* aged, **b**) and data type (scRNA-seq *vs.* snRNA-seq, **c**). **d**, **f**, **g,** UMAP plot illustrating 96,529 cells/nuclei from mouse skeletal muscle across age with major cell types(**d**), age group (young *vs.* aged, **f**) and data type (scRNA-seq *vs.* snRNA-seq, **g**) shown. **e**, Dot plot of marker genes for each cell type in mouse skeletal muscle ageing atlas. Size of the dot represents the proportion of cells expressing a gene. Colour denotes scaled expression level.

**Extended Data Fig. 2.**
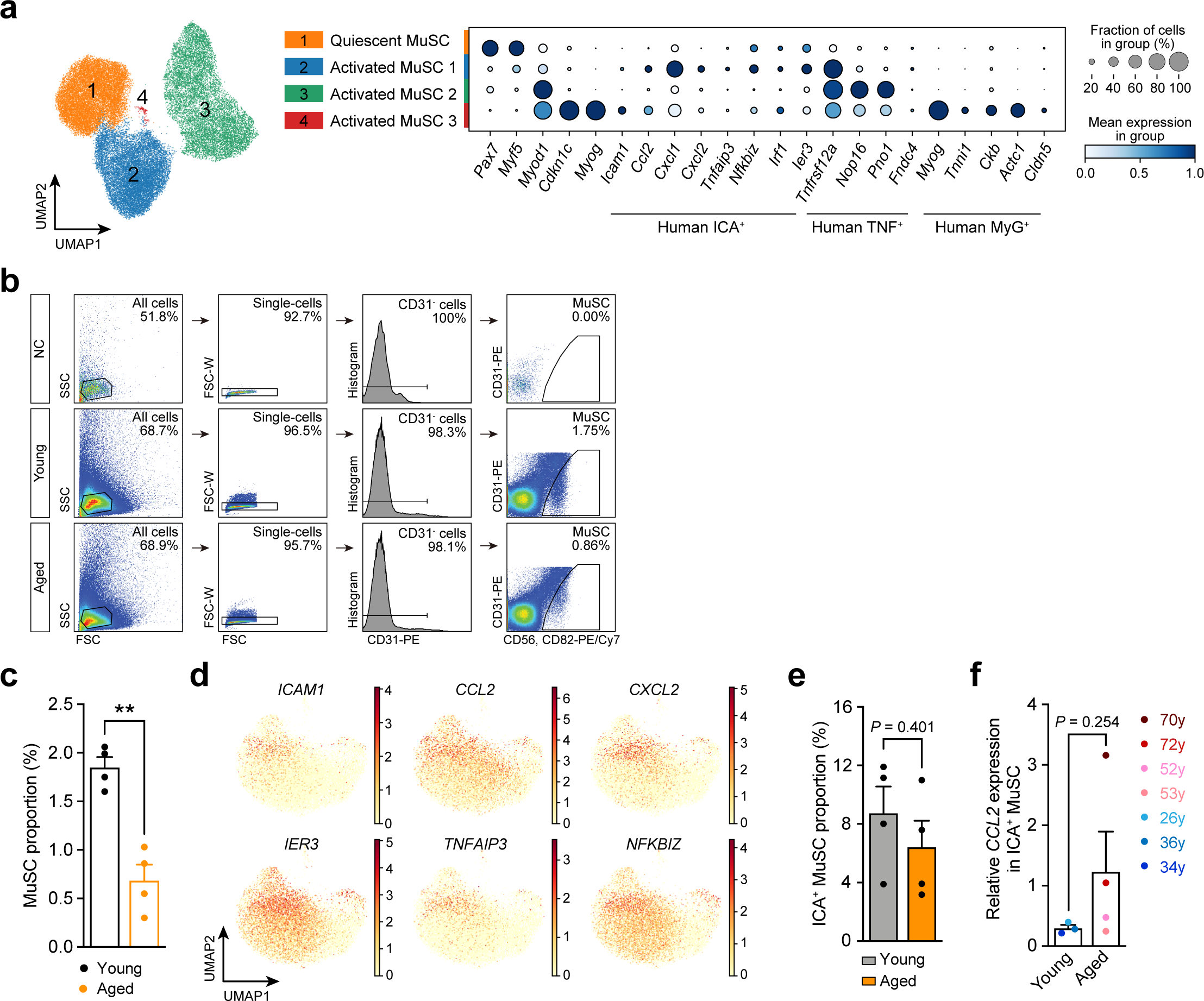
Mechanistic insights into human MuSC ageing. **a**, UMAP plot of mouse MuSC subpopulations obtained via integration of 5 publicly available mouse resources, annotation into quiescent and activated populations was borrowed from corresponding datasets (see Methods). Marker genes corresponding to human MuSC subpopulations are indicated. Size of the dot represents the proportion of cells expressing a gene. Colour denotes scaled expression level. **b**, Gating strategy for FACS-based MuSC sorting. **c**, Bar plot showing change in the proportion of FACS-sorted human MuSC, relative to all isolated cells, with age (n = 4 for young *vs.* n = 4 for aged). *P* value was obtained using unpaired two-tailed *t*-test. **, *P* < 0.01. **d**, UMAP plots showing expression of immune-related genes enriched in ICA^+^ MuSC including *CCL2*, *CXCL2*, *IER3*, *TNFAIP3* and *NFKBIZ*. **e**, Bar plot showing proportion of ICA^+^ MuSC, relative to the total MuSC, in young *vs.* aged (n = 4 for young *vs.* n = 4 for aged). *P* value was obtained using unpaired two-tailed *t*-test. **f**, Bar plots showing change in the relative expression of *CCL2*, quantified with real-time PCR, in ICA^+^ MuSC during human ageing (n = 3 for young *vs.* n = 4 for aged). *P* value was obtained using unpaired *t*-test with Welch’s correction two-tailed test. All data presented in bar plots (**c**, **e**, **f**) are expressed as mean ± SE with individual data points shown.

**Extended Data Fig. 3.**
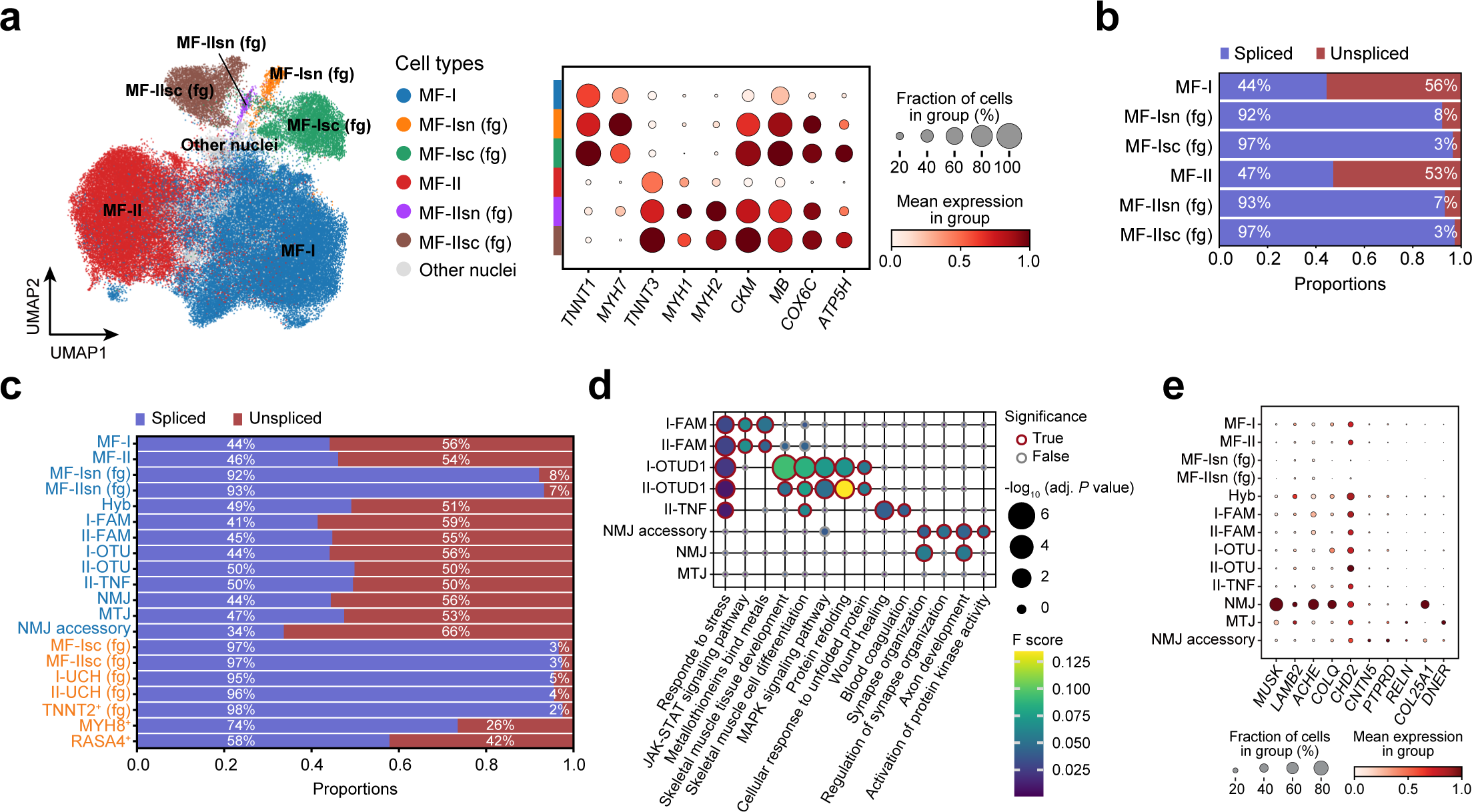
Myofiber type-specific ageing mechanisms. **a**, UMAP visualisation of myofiber populations obtained from integrated sn- and scRNA-seq dataset, coloured according to the six main populations (left) and dot plot showing main marker genes (right). Size of the dot represents the proportion of cells expressing a gene. Colour denotes scaled expression level **b**, **c**, Bar plots illustrating percentage of spliced and unspliced transcripts relative to the total for the main (**b**) and subtype (**c**) myofiber populations. **d**, Dot plot showing selected gene sets and their enrichment in the I and II-FAM, I and II-OTU, II-TNF and NMJ accessory populations based on gProfiler over-representation analysis using populations’ marker genes (see Methods). Colour denotes F score. Size of the dot represents -log_10_ of adjusted (adj.) *P* value, significant values are marked with a red edge. **e**, Dot plot showing classical NMJ markers which were rarely expressed in NMJ accessory nuclei as well as synapse-related genes specific to NMJ accessory nuclei. Size of the dot represents the proportion of cells expressing a gene. Colour denotes scaled expression level.

**Extended Data Fig. 4.**
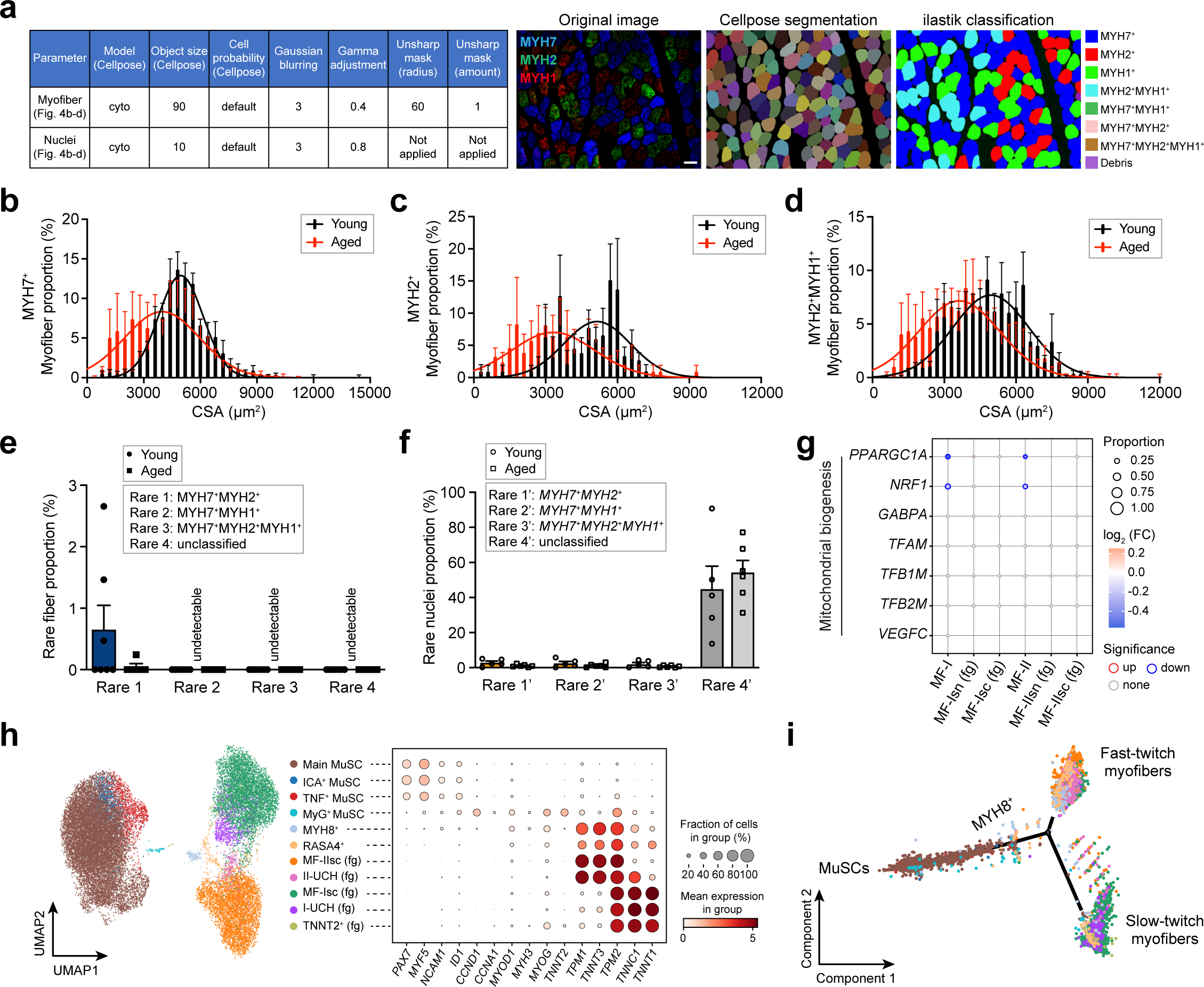
Compensatory mechanisms against human fast-twitch myofiber degeneration in ageing. **a**, Table of parameters used for Cellpose myofiber and nuclei segmentation (left) and example images from different stages of image analysis workflow (right). **b**-**d**, Histogram illustrating distribution of the myofiber cross-sectional area (CSA, x axis) and its change with age (n=7 for young *vs.* n=5 for aged) for MYH7^+^ (**b**), MYH2^+^ (**c**) or MYH2^+^MYH1^+^ (**d**) myofiber, respectively. Gaussian curve fits were obtained using the nonlinear regression test. **e**, **f**, Paired bar plots showing proportion of rare or unclassified myofiber (**e**) and myonuclei (**f**) types in young *vs.* aged individuals. **g**, Dot plot illustrating age-associated changes in the mitochondrial biogenesis genes for six main myofiber populations. Size of the dot represents the proportion of aged cells expressing the gene. Colour denotes log_2_ (FC) in gene expression in aged *vs.* young individuals. Significantly up-regulated genes are highlighted with red and down-regulated with blue edges. **h**, UMAP plot (left) shows MuSCs and myofiber populations from scRNA-seq. Dot plot (right) shows their marker genes, which are presented in the order of their appearance in the myogenesis trajectory. Size of the dot represents the proportion of cells expressing a gene. Colour denotes scaled expression level. **i**, Reduce dimensional space showing cellular trajectory inferred by Monocle2 algorithm between MuSC and myofiber. All data presented in bar plots (**b**-**f**) are expressed as mean ± SE with individual data points shown.

**Extended Data Fig. 5.**
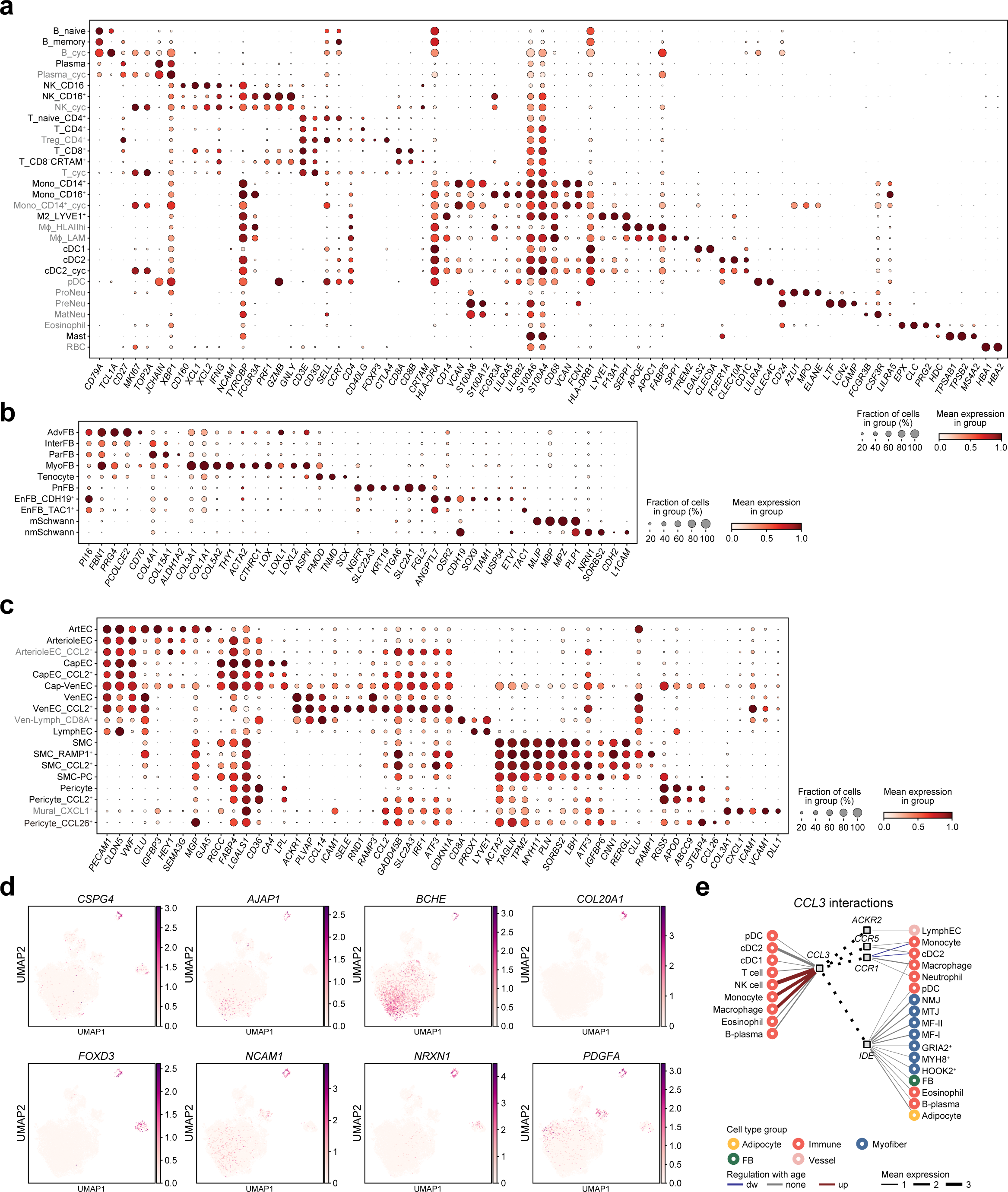
Cell type composition of human skeletal muscle microenvironment and ageing regulation. **a**-**c**, Dot plots illustrating marker genes specific for subpopulations of immune cells (**a**), fibroblasts and Schwann cells (**b**) as well as endothelial and smooth muscle cells (**c**). Size of the dot represents the proportion of cells expressing a gene. Colour denotes scaled expression level. **d**, UMAP plot showing specificity of nonmyelinating Schwann cells characterised by expression of *CSPG4*, *AJAP1*, *BCHE*, *COL20A1*, *FOXD3*, *NCAM1*, *NRXN1* and *PDGFA*. **e**, Putative cell-cell interactions in the aged skeletal muscle mediated via *CCL3* chemokine produced by microenvironment cells (see Fig. 5f).

**Extended Data Fig. 6.**
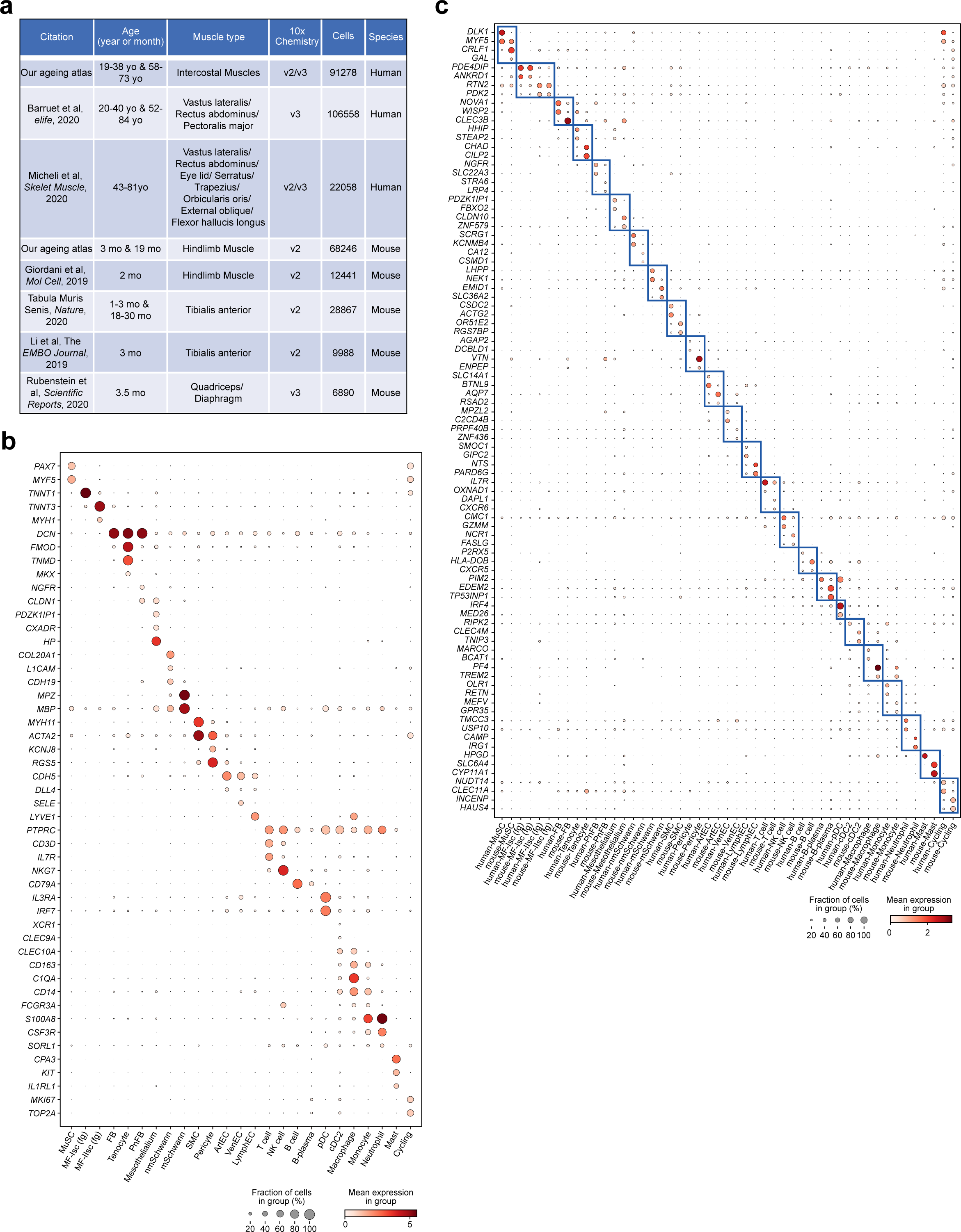
Integrated human-mouse skeletal muscle ageing atlas at single-cell resolution. **a**, Overview of the publicly available datasets used to make the human-mouse skeletal muscle ageing atlas. **b**, **c**, Dot plot showing species-common (**b**) and -specific (**c**) marker genes for each major cell type annotated in the human-mouse skeletal muscle ageing atlas. Size of the dot represents the proportion of cells expressing a gene. Colour denotes scaled expression level.

## References

1. López-Otín, C., Blasco, M. A., Partridge, L., Serrano, M. & Kroemer, G. The hallmarks of aging. Cell 153, 1194–1217 (2013).

2. Huard, J., Li, Y. & Fu, F. H. Muscle injuries and repair: current trends in research. J. Bone Joint Surg. Am. 84, 822–832 (2002).

3. Pedersen, B. K. & Febbraio, M. A. Muscles, exercise and obesity: skeletal muscle as a secretory organ. Nat. Rev. Endocrinol. 8, 457–465 (2012).

4. Tidball, J. G. Regulation of muscle growth and regeneration by the immune system. Nat. Rev. Immunol. 17, 165–178 (2017).

5. Hargreaves, M. & Spriet, L. L. Skeletal muscle energy metabolism during exercise. Nat Metab 2, 817–828 (2020).

6. Siparsky, P. N., Kirkendall, D. T. & Garrett, W. E., Jr. Muscle changes in aging: understanding sarcopenia. Sports Health 6, 36–40 (2014).

7. Cruz-Jentoft, A. J. & Sayer, A. A. Sarcopenia. Lancet 393, 2636–2646 (2019).

8. Yeung, S. S. Y. et al. Sarcopenia and its association with falls and fractures in older adults: A systematic review and meta-analysis. J. Cachexia Sarcopenia Muscle 10, 485–500 (2019).

9. Falls. https://www.who.int/news-room/fact-sheets/detail/falls.

10. Nilwik, R. et al. The decline in skeletal muscle mass with aging is mainly attributed to a reduction in type II muscle fiber size. Exp. Gerontol. 48, 492–498 (2013).

11. Gopinath, S. D. & Rando, T. A. Stem cell review series: aging of the skeletal muscle stem cell niche. Aging Cell 7, 590–598 (2008).

12. Zhang, H. et al. NAD^+^ repletion improves mitochondrial and stem cell function and enhances life span in mice. Science 352, 1436–1443 (2016).

13. Schiffer, I. et al. miR-1 coordinately regulates lysosomal v-ATPase and biogenesis to impact proteotoxicity and muscle function during aging. Elife 10, (2021).

14. Chini, C. C. S. et al. CD38 ecto-enzyme in immune cells is induced during aging and regulates NAD+ and NMN levels. Nat Metab 2, 1284–1304 (2020).

15. Heredia, J. E. et al. Type 2 innate signals stimulate fibro/adipogenic progenitors to facilitate muscle regeneration. Cell 153, 376–388 (2013).

16. Kuswanto, W. et al. Poor Repair of Skeletal Muscle in Aging Mice Reflects a Defect in Local, Interleukin-33-Dependent Accumulation of Regulatory T Cells. Immunity 44, 355–367 (2016).

17. McKellar, D. W. et al. Large-scale integration of single-cell transcriptomic data captures transitional progenitor states in mouse skeletal muscle regeneration. Commun Biol 4, 1280 (2021).

18. De Micheli, A. J., Spector, J. A., Elemento, O. & Cosgrove, B. D. A reference single-cell transcriptomic atlas of human skeletal muscle tissue reveals bifurcated muscle stem cell populations. Skelet. Muscle 10, 19 (2020).

19. Rubenstein, A. B. et al. Single-cell transcriptional profiles in human skeletal muscle. Sci. Rep. 10, 229 (2020).

20. De Micheli, A. J. et al. Single-Cell Analysis of the Muscle Stem Cell Hierarchy Identifies Heterotypic Communication Signals Involved in Skeletal Muscle Regeneration. Cell Rep. 30, 3583–3595.e5 (2020).

21. Barruet, E. et al. Functionally heterogeneous human satellite cells identified by single cell RNA sequencing. Elife 9, (2020).

22. Kim, M. et al. Single-nucleus transcriptomics reveals functional compartmentalization in syncytial skeletal muscle cells. Nat. Commun. 11, 6375 (2020).

23. Dos Santos, M. et al. Single-nucleus RNA-seq and FISH identify coordinated transcriptional activity in mammalian myofibers. Nat. Commun. 11, 5102 (2020).

24. Petrany, M. J. et al. Single-nucleus RNA-seq identifies transcriptional heterogeneity in multinucleated skeletal myofibers. Nat. Commun. 11, 6374 (2020).

25. Perez, K. et al. Single nuclei profiling identifies cell specific markers of skeletal muscle aging, sarcopenia and senescence. bioRxiv (2021) doi:10.1101/2021.01.22.21250336.

26. Orchard, P. et al. Human and rat skeletal muscle single-nuclei multi-omic integrative analyses nominate causal cell types, regulatory elements, and SNPs for complex traits. Genome Res. (2021) doi:10.1101/gr.268482.120.

27. Gayoso, A. et al. scvi-tools: a library for deep probabilistic analysis of single-cell omics data. bioRxiv 2021.04.28.441833 (2021) doi:10.1101/2021.04.28.441833.

28. Dulken, B. W. et al. Single-cell analysis reveals T cell infiltration in old neurogenic niches. Nature 571, 205–210 (2019).

29. Schaum, N. et al. Ageing hallmarks exhibit organ-specific temporal signatures. Nature 583, 596–602 (2020).

30. Sousa-Victor, P., García-Prat, L. & Muñoz-Cánoves, P. Control of satellite cell function in muscle regeneration and its disruption in ageing. Nat. Rev. Mol. Cell Biol. 23, 204–226 (2022).

31. Relaix, F. et al. Perspectives on skeletal muscle stem cells. Nat. Commun. 12, 692 (2021).

32. Tierney, M. T., Stec, M. J., Rulands, S., Simons, B. D. & Sacco, A. Muscle Stem Cells Exhibit Distinct Clonal Dynamics in Response to Tissue Repair and Homeostatic Aging. Cell Stem Cell 22, 119–127.e3 (2018).

33. Saber, J., Lin, A. Y. T. & Rudnicki, M. A. Single-cell analyses uncover granularity of muscle stem cells. F1000Rs. 9, (2020).

34. Novak, J. S. et al. Human muscle stem cells are refractory to aging. Aging Cell 20, e13411 (2021).

35. Carlson, M. E. et al. Molecular aging and rejuvenation of human muscle stem cells. EMBO Mol. Med. 1, 381–391 (2009).

36. Sharifi, S., da Costa, H. F. R. & Bierhoff, H. The circuitry between ribosome biogenesis and translation in stem cell function and ageing. Mech. Ageing Dev. 189, 111282 (2020).

37. Giordani, L. et al. High-Dimensional Single-Cell Cartography Reveals Novel Skeletal Muscle-Resident Cell Populations. Mol. Cell 74, 609–621.e6 (2019).

38. Kimmel, J. C., Hwang, A. B., Scaramozza, A., Marshall, W. F. & Brack, A. S. Aging induces aberrant state transition kinetics in murine muscle stem cells. Development 147, (2020).

39. Li, H. et al. Muscle-secreted granulocyte colony-stimulating factor functions as metabolic niche factor ameliorating loss of muscle stem cells in aged mice. EMBO J. 38, e102154 (2019).

40. Tabula Muris Consortium. A single-cell transcriptomic atlas characterizes ageing tissues in the mouse. Nature 583, 590–595 (2020).

41. Dann, E., Henderson, N. C., Teichmann, S. A., Morgan, M. D. & Marioni, J. C. Differential abundance testing on single-cell data using k-nearest neighbor graphs. Nat. Biotechnol. 40, 245–253 (2022).

42. Alexander, M. S. et al. CD82 Is a Marker for Prospective Isolation of Human Muscle Satellite Cells and Is Linked to Muscular Dystrophies. Cell Stem Cell 19, 800–807 (2016).

43. Magli, A. et al. PAX7 Targets, CD54, Integrin α9β1, and SDC2, Allow Isolation of Human ESC/iPSC-Derived Myogenic Progenitors. Cell Rep. 19, 2867–2877 (2017).

44. Lessard, F. et al. Senescence-associated ribosome biogenesis defects contributes to cell cycle arrest through the Rb pathway. Nat. Cell Biol. 20, 789–799 (2018).

45. Nishimura, K. et al. Perturbation of ribosome biogenesis drives cells into senescence through 5S RNP-mediated p53 activation. Cell Rep. 10, 1310–1323 (2015).

46. Ziemkiewicz, N., Hilliard, G., Pullen, N. A. & Garg, K. The Role of Innate and Adaptive Immune Cells in Skeletal Muscle Regeneration. Int. J. Mol. Sci. 22, (2021).

47. Bengal, E., Perdiguero, E., Serrano, A. L. & Muñoz-Cánoves, P. Rejuvenating stem cells to restore muscle regeneration in aging. F1000Res. 6, 76 (2017).

48. Thompson, W. L. & Van Eldik, L. J. Inflammatory cytokines stimulate the chemokines CCL2/MCP-1 and CCL7/MCP-3 through NFkB and MAPK dependent pathways in rat astrocytes [corrected]. Brain Res. 1287, 47–57 (2009).

49. Rogala, K. B. et al. Structural basis for the docking of mTORC1 on the lysosomal surface. Science 366, 468–475 (2019).

50. Petit, C. S., Roczniak-Ferguson, A. & Ferguson, S. M. Recruitment of folliculin to lysosomes supports the amino acid-dependent activation of Rag GTPases. J. Cell Biol. 202, 1107–1122 (2013).

51. Tsun, Z.-Y. et al. The folliculin tumor suppressor is a GAP for the RagC/D GTPases that signal amino acid levels to mTORC1. Mol. Cell 52, 495–505 (2013).

52. Frederick, D. W. et al. Loss of NAD Homeostasis Leads to Progressive and Reversible Degeneration of Skeletal Muscle. Cell Metab. 24, 269–282 (2016).

53. Basse, A. L. et al. Nampt controls skeletal muscle development by maintaining Ca2+ homeostasis and mitochondrial integrity. Mol Metab 53, 101271 (2021).

54. Moresi, V., Adamo, S. & Berghella, L. The JAK/STAT Pathway in Skeletal Muscle Pathophysiology. Front. Physiol. 10, 500 (2019).

55. Caldow, M. K., Steinberg, G. R. & Cameron-Smith, D. Impact of SOCS3 overexpression on human skeletal muscle development in vitro. Cytokine 55, 104–109 (2011).

56. Serrano, A. L., Baeza-Raja, B., Perdiguero, E., Jardí, M. & Muñoz-Cánoves, P. Interleukin-6 is an essential regulator of satellite cell-mediated skeletal muscle hypertrophy. Cell Metab. 7, 33–44 (2008).

57. Sun, L. et al. JAK1-STAT1-STAT3, a key pathway promoting proliferation and preventing premature differentiation of myoblasts. J. Cell Biol. 179, 129–138 (2007).

58. Doles, J. D. & Olwin, B. B. The impact of JAK-STAT signaling on muscle regeneration. Nat. Med. 20, 1094–1095 (2014).

59. Price, F. D. et al. Inhibition of JAK-STAT signaling stimulates adult satellite cell function. Nat. Med. 20, 1174–1181 (2014).

60. Tierney, M. T. et al. STAT3 signaling controls satellite cell expansion and skeletal muscle repair. Nat. Med. 20, 1182–1186 (2014).

61. Sato, S., Ogura, Y. & Kumar, A. TWEAK/Fn14 Signaling Axis Mediates Skeletal Muscle Atrophy and Metabolic Dysfunction. Front. Immunol. 5, 18 (2014).

62. Pascoe, A. L., Johnston, A. J. & Murphy, R. M. Controversies in TWEAK-Fn14 signaling in skeletal muscle atrophy and regeneration. Cell. Mol. Life Sci. 77, 3369–3381 (2020).

63. Ramachandran, P. et al. Resolving the fibrotic niche of human liver cirrhosis at single-cell level. Nature 575, 512–518 (2019).

64. Colombo, M. N. & Francolini, M. Glutamate at the Vertebrate Neuromuscular Junction: From Modulation to Neurotransmission. Cells 8, (2019).

65. Henley, J. M. & Wilkinson, K. A. Synaptic AMPA receptor composition in development, plasticity and disease. Nat. Rev. Neurosci. 17, 337–350 (2016).

66. Bonanomi, D. et al. Ret is a multifunctional coreceptor that integrates diffusible- and contact-axon guidance signals. Cell 148, 568–582 (2012).

67. Hallock, P. T., Chin, S., Blais, S., Neubert, T. A. & Glass, D. J. Sorbs1 and -2 Interact with CrkL and Are Required for Acetylcholine Receptor Cluster Formation. Mol. Cell. Biol. 36, 262–270 (2016).

68. Murgia, M. et al. Single Muscle Fiber Proteomics Reveals Fiber-Type-Specific Features of Human Muscle Aging. Cell Rep. 19, 2396–2409 (2017).

69. Schiaffino, S., Rossi, A. C., Smerdu, V., Leinwand, L. A. & Reggiani, C. Developmental myosins: expression patterns and functional significance. Skelet. Muscle 5, 22 (2015).

70. Evano, B. & Tajbakhsh, S. Skeletal muscle stem cells in comfort and stress. NPJ Regen Med 3, 24 (2018).

71. Conde, C. D. et al. Cross-tissue immune cell analysis reveals tissue-specific adaptations and clonal architecture across the human body. bioRxiv 2021.04.28.441762 (2021) doi:10.1101/2021.04.28.441762.

72. Chakarov, S. et al. Two distinct interstitial macrophage populations coexist across tissues in specific subtissular niches. Science 363, (2019).

73. Cui, C.-Y. & Ferrucci, L. Macrophages in skeletal muscle aging. Aging 12, 3–4 (2020).

74. Eraslan, G. et al. Single-nucleus cross-tissue molecular reference maps to decipher disease gene function. bioRxiv 2021.07.19.452954 (2021) doi:10.1101/2021.07.19.452954.

75. Soares, M. P. & Hamza, I. Macrophages and Iron Metabolism. Immunity 44, 492–504 (2016).

76. Jaitin, D. A. et al. Lipid-Associated Macrophages Control Metabolic Homeostasis in a Trem2-Dependent Manner. Cell 178, 686–698.e14 (2019).

77. Forcina, L., Cosentino, M. & Musarò, A. Mechanisms Regulating Muscle Regeneration: Insights into the Interrelated and Time-Dependent Phases of Tissue Healing. Cells 9, (2020).

78. Buechler, M. B. et al. Cross-tissue organization of the fibroblast lineage. Nature 593, 575–579 (2021).

79. Castro, R. et al. Specific labeling of synaptic schwann cells reveals unique cellular and molecular features. Elife 9, (2020).

80. Madissoon, E. et al. A spatial multi-omics atlas of the human lung reveals a novel immune cell survival niche. bioRxiv 2021.11.26.470108 (2021) doi:10.1101/2021.11.26.470108.

81. Chen, R. et al. CD147 deficiency in T cells prevents thymic involution by inhibiting the EMT process in TECs in the presence of TGFβ. Cell. Mol. Immunol. 18, 171–181 (2021).

82. Jensen, G. L. Inflammation: roles in aging and sarcopenia. JPEN J. Parenter. Enteral Nutr. 32, 656–659 (2008).

83. Karanth, S. D. et al. Inflammation in Relation to Sarcopenia and Sarcopenic Obesity among Older Adults Living with Chronic Comorbidities: Results from the National Health and Nutrition Examination Survey 1999-2006. Nutrients 13, (2021).

84. Fukada, K. & Kajiya, K. Age-related structural alterations of skeletal muscles and associated capillaries. Angiogenesis 23, 79–82 (2020).

85. Efremova, M., Vento-Tormo, M., Teichmann, S. A. & Vento-Tormo, R. CellPhoneDB: inferring cell-cell communication from combined expression of multi-subunit ligand-receptor complexes. Nat. Protoc. 15, 1484–1506 (2020).

86. Lu, H., Huang, D., Ransohoff, R. M. & Zhou, L. Acute skeletal muscle injury: CCL2 expression by both monocytes and injured muscle is required for repair. FASEB J. 25, 3344–3355 (2011).

87. Warren, G. L. et al. Role of CC chemokines in skeletal muscle functional restoration after injury. Am. J. Physiol. Cell Physiol. 286, C1031–6 (2004).

88. Hirata, A. et al. Expression profiling of cytokines and related genes in regenerating skeletal muscle after cardiotoxin injection: a role for osteopontin. Am. J. Pathol. 163, 203–215 (2003).

89. Tanaka, T., Narazaki, M. & Kishimoto, T. IL-6 in inflammation, immunity, and disease. Cold Spring Harb. Perspect. Biol. 6, a016295 (2014).

90. Howes, A., Gabryšová, L. & O’Garra, A. Role of IL-10 and the IL-10 Receptor in Immune Responses. in Reference Module in Biomedical Sciences (Elsevier, 2014).

91. Zhuang, J. et al. Comparison of multi-tissue aging between human and mouse. Sci. Rep. 9, 6220 (2019).

92. Zhang, M. J., Pisco, A. O., Darmanis, S. & Zou, J. Mouse aging cell atlas analysis reveals global and cell type-specific aging signatures. Elife 10, (2021).

93. Vyas, J. M., Van der Veen, A. G. & Ploegh, H. L. The known unknowns of antigen processing and presentation. Nat. Rev. Immunol. 8, 607–618 (2008).

94. Oikonomopoulou, K., Ricklin, D., Ward, P. A. & Lambris, J. D. Interactions between coagulation and complement--their role in inflammation. Semin. Immunopathol. 34, 151–165 (2012).

95. Enge, M. et al. Single-Cell Analysis of Human Pancreas Reveals Transcriptional Signatures of Aging and Somatic Mutation Patterns. Cell 171, 321–330.e14 (2017).

96. Zou, Z. et al. A Single-Cell Transcriptomic Atlas of Human Skin Aging. Dev. Cell 56, 383–397.e8 (2021).

97. Yi, W. et al. A single-cell transcriptome atlas of the aging human and macaque retina. Natl Sci Rev 8, nwaa179 (2021).

98. Webster, C., Silberstein, L., Hays, A. P. & Blau, H. M. Fast muscle fibers are preferentially affected in Duchenne muscular dystrophy. Cell 52, 503–513 (1988).

99. Zhang, H. Single cell isolation from human skeletal muscle v1 (protocols.io.q5wdy7e). doi:10.17504/protocols.io.q5wdy7e.

100. Zhang, H. Nuclei isolation from human skeletal muscle v1 (protocols.io.t68erhw). doi:10.17504/protocols.io.t68erhw.

101. Kaminow, B., Yunusov, D. & Dobin, A. STARsolo: accurate, fast and versatile mapping/quantification of single-cell and single-nucleus RNA-seq data. bioRxiv 2021.05.05.442755 (2021) doi:10.1101/2021.05.05.442755.

102. Fleming, S. J., Marioni, J. C. & Babadi, M. CellBender remove-background: a deep generative model for unsupervised removal of background noise from scRNA-seq datasets. bioRxiv 791699 (2019) doi:10.1101/791699.

103. Wolock, S. L., Lopez, R. & Klein, A. M. Scrublet: Computational Identification of Cell Doublets in Single-Cell Transcriptomic Data. Cell Syst 8, 281–291.e9 (2019).

104. Wolf, F. A., Angerer, P. & Theis, F. J. SCANPY: large-scale single-cell gene expression data analysis. Genome Biol. 19, 15 (2018).

105. Raudvere, U. et al. g:Profiler: a web server for functional enrichment analysis and conversions of gene lists (2019 update). Nucleic Acids Res. 47, W191–W198 (2019).

106. Zhou, Y. et al. Metascape provides a biologist-oriented resource for the analysis of systems-level datasets. Nat. Commun. 10, 1523 (2019).

107. Trapnell, C. et al. The dynamics and regulators of cell fate decisions are revealed by pseudotemporal ordering of single cells. Nat. Biotechnol. 32, 381–386 (2014).

108. Yoshida, M. et al. Local and systemic responses to SARS-CoV-2 infection in children and adults. Nature 602, 321–327 (2022).

109. Young, A. M. H. et al. A map of transcriptional heterogeneity and regulatory variation in human microglia. Nat. Genet. 53, 861–868 (2021).

110. van der Walt, S. et al. scikit-image: image processing in Python. PeerJ 2, e453 (2014).

111. Stringer, C., Wang, T., Michaelos, M. & Pachitariu, M. Cellpose: a generalist algorithm for cellular segmentation. Nat. Methods 18, 100–106 (2021).

112. Berg, S. et al. ilastik: interactive machine learning for (bio)image analysis. Nat. Methods 16, 1226–1232 (2019).

113. Kummer, T. T., Misgeld, T., Lichtman, J. W. & Sanes, J. R. Nerve-independent formation of a topologically complex postsynaptic apparatus. J. Cell Biol. 164, 1077–1087 (2004).

114. Nishimune, H. et al. Laminins promote postsynaptic maturation by an autocrine mechanism at the neuromuscular junction. J. Cell Biol. 182, 1201–1215 (2008).

