## Supplementary Tables for "Human skeletal muscle ageing atlas"

Legends for all supplementary tables:
 

Supplementary Table 1:
(1) Table containing the available donor metadata; (2,3) Table of the top 100 marker genes for the major cell populations identified in the human (2) and mouse (3) integrated single-cell and single-nuclei ageing cell atlases. Marker genes were calculated using t-test, by comparing “cluster” of interest vs. the rest and P values were corrected for multiple comparisons using Benjamini-Hochberg correction.

Supplementary Table 2: 
(1) Top 100 marker genes for MuSCs subtypes identified using t-test followed by  Benjamini-Hochberg correction; (2) Gene Ontology enrichment results for the markers of MuSCs cell subtypes obtained using Metascape web service. Only top 10 clusters per each MuSC population and top 5 enriched terms within each cluster were selected. (3) Ribosome biogenesis enrichment scores for MuSC subtypes within each donor.

Supplementary Table 3: 
(1,2) Ageing differential expression results for every major cell type (1) and every subtype (2) in the integrated single-cell and single-nuclei human ageing cell atlas. Effect sizes were derived from linear mixed-effects regression model, accounting for the technical (data modality, 10X chemistry) and biological (donor, sex) covariates;
(3) Ageing differential gene expression results for every cell subtype in single-cell  human ageing cell atlas derived from linear mixed-effects regression model, accounting for the technical (data modality, 10X chemistry) and biological (donor, sex) covariates.

Supplementary Table 4: 
(1,2) Table of the marker genes for the subpopulations identified in human myofiber snRNAseq (1) and scRNAseq (2). The genes were obtained using scVI gene expression model while testing current population vs. the rest, genes were prefiltered to be expressed in at least 10 % of cells, to have log2 (Fold change, FC) above 1 and bayes factor above 2. (3) Table of the marker genes for selected myonuclei populations (I and II-FAM+, I and OTU+, II-TNF and NMJ accessory) obtained by differential expression between the mentioned populations and baseline type I and type II myonuclei clusters. Genes were prefiltered to be expressed in at least 10 % of cells, to have log2 (Fold change, FC) above 1 and bayes factor above 2. (4) Gene set enrichment results for selected myonuclei populations (I and II-FAM+, I and OTU+, II-TNF and NMJ accessory) performed using gProfiler. Genes in (3) were used as an input, GO BP, KEGG and Reactome gene sets were used for enrichment. 

Supplementary Table 5: 
(1) Table output from cellpose segmentation, followed by ilastik classification, on images from MYH1+, MYH2+, MYH7+ immunohistochemistry (IHC) stained tissue sections; (2) Myofiber type count for each donor from IHC. (3) Myonuclei type count for each donor from snRNAseq; (4,5) Quantification of MYH8+ expression from scRNAseq (4) and from immunofluorescence (IF) (5).(6) Quantification of centralised nuclei and their co-expression with fast-twitch MYH2+ myofibers from IF.


Supplementary Table 6: 
(1,2,3) Table of the top 100 marker genes for the cell subpopulations identified in
the immune (1), fibroblast and Schwann (2) and blood vessel (3) compartments.  and mouse (3) integrated single-cell and single-nuclei ageing cell atlas. Marker genes were calculated using t-test, by comparing “cluster” of interest vs. the rest and P values were corrected for multiple comparisons using Benjamini-Hochberg correction;(4) The list of putative cell-cell interactions pairs mediated via CCL2, CCL3, CCL4 and CXCL8 ligands which were obtained using CellphoneDB resource;

Supplementary Table 7: 

(1,2) Top 100 common (1) and species-specific (2) marker genes for the major cell types identified in the integrated human-mouse single-cell ageing dataset. Marker genes were calculated using t-test, by comparing “cluster” of interest vs. the rest and P values were corrected for multiple comparisons using Benjamini-Hochberg correction; (3,4) Ageing differential expression results for the major human (3) and mouse (4) cell types within the integrated human-mouse single-cell ageing dataset. Effect sizes were derived from linear mixed-effects regression model, accounting for the technical (10X chemistry, data source) and biological (donor, muscle type, sex) covariates; (5) Table of KEGG pathways displaying enrichment in ageing DEGs in the same cell type both in mouse and human according to ClusterProfiler. 
